# Spatial immune profiling reveals lymphocyte confinement to myeloid–mesenchymal niches and heterogeneous therapeutic T-cell opportunity in pediatric ependymoma

**DOI:** 10.64898/2026.09.24.754209

**Authors:** Patrick Truong, Reza Mirzazadeh, Joakim Lundeberg, Klas Blomgren, Lola Boutin

## Abstract

Pediatric posterior fossa group A (PFA) ependymoma is an aggressive, chemo-resistant brain tumor with a poor 10-year overall survival and no therapeutic options at relapse. Although conventionally classified as immune-cold, the spatial organization and therapeutic relevance of its rare immune infiltrates remain poorly characterized. Here, we present a spatially resolved immune analysis of 14 PFA ependymoma sections using published Visium spatial transcriptomics data analyzed through a multi-stage immune-focused computational framework combining reference-guided deconvolution, functional lymphocyte state scoring, ligand–receptor inference, and αβ/γδ T cell immunotherapy opportunity scoring.

Lymphocyte-associated hotspots were identified in 6 of 14 sections and, among confidently classified hotspots, were dominated by inflammatory-associated and retention-associated transcriptional states rather than cytotoxic or exhausted programs. These hotspots were preferentially confined to myeloid- and mesenchymal-rich tumor zones with near-complete exclusion from epithelial regions; ligand–receptor analysis identified four candidate restriction axes (SPP1–CD44, FN1–integrin, collagen VI–integrin, and APP–CD74). Immunotherapy opportunity scoring revealed heterogeneous and spatially compartmentalized αβ and γδ T cell recognition landscapes across sections: αβT ligand availability and immunosuppressive tone both rose in the mesenchymal compartment, while the two components of γδ phosphoantigen-mediated recognition were spatially and transcriptionally dissociated from one another.

Together, these findings reframe PFA ependymoma not as a monolithic immune desert but as a tumor type with spatially compartmentalized, mechanistically tractable immune niches, and provide a generalizable computational strategy for mapping immune architecture and T cell therapy susceptibility in low-infiltrated pediatric CNS tumors.

## Introduction

Ependymal tumors are neuroepithelial malignancies that arise from the ependymal lining of the cerebral ventricles and central canal of the spinal cord. They occur across all age groups but disproportionately affect children^1^. In the pediatric setting, approximately 90% of ependymomas arise intracranially, with roughly two-thirds localized to the posterior fossa (PF) and one-third to the supratentorial (ST) compartment^2^. Despite multimodal treatment combining maximal surgical resection and conformal radiotherapy, long-term outcomes remain unsatisfactory. Population-level data from the National Cancer Database, encompassing over 900 pediatric patients diagnosed between 2010 and 2017, reported overall survival (OS) rates of 89%, 82.9%, and 74.5% at three, five, and ten years, respectively^3^. Outcomes are particularly poor in infants, with a five-year OS reaching only 67% in dedicated infant cohorts, where the extent of resection and chemotherapy exposure emerge as the primary independent prognostic factors^4^. Critically, no effective systemic salvage therapy has been established for relapsed disease, underscoring the persistent and urgent unmet clinical need.

Genome-wide DNA methylation profiling has resolved nine biologically and clinically distinct molecular subgroups of ependymoma distributed across three CNS compartments^5^, a classification formally incorporated into the 2021 WHO classification of CNS tumors^6^. Within the posterior fossa (PF), two subgroups have been defined with strikingly divergent outcomes: PF group B (PFB), which predominantly affects older children and adolescents and carries an excellent prognosis with no observed relapses in molecularly stratified cohorts, and PF group A (PFA), which predominantly affects young children and represents the most common and lethal subtype^5^. Long-term outcome data from molecularly defined cohorts confirm the dismal prognosis of PFA, with a five-year progression-free survival (PFS) of approximately 43% and a ten-year PFS below 40%^7^. The biological drivers underlying this poor outcome and the absence of targetable therapeutic vulnerabilities remain incompletely understood, motivating deeper investigation of the PFA tumor microenvironment (TME).

Pediatric brain tumors are broadly regarded as immunologically “cold,” characterized by a low mutational burden, scarce neoantigen load, and limited expression of immune checkpoint ligands such as PD-L1^8^. Consistently, large-scale transcriptional immunogenomic profiling of 925 pediatric nervous system tumors spanning 12 histological types has resolved four distinct immune phenotypes: Pediatric Inflamed, Myeloid Predominant, Immune Neutral, and Immune Desert, with ependymomas disproportionately represented in the less-inflamed clusters, reinforcing its classification as an immune-restricted tumor type^9^. Pan-histological analyses of shared immunosuppressive programs have identified common molecular vulnerabilities across these clusters that may be amenable to targeted immunomodulation^10^. Nevertheless, the therapeutic pressure of chemoresistance and poor outcomes have driven growing interest in immunotherapeutic strategies for pediatric brain tumors, encompassing checkpoint inhibition, cancer vaccines, and adoptive cellular therapies, including CAR-T cells, each of which faces distinct barriers in the pediatric CNS context, ranging from limited target antigen expression to an immunosuppressive TME and restricted immune cell trafficking across the blood-brain barrier^11^. Identifying which tumors harbor focal immune susceptibility and the microenvironmental programs that govern it is a prerequisite for rational immunotherapy design. Increasing evidence suggests that the immune landscape of pediatric CNS tumors is heterogeneous and biologically informative, despite low overall infiltration. Large-scale methylation-based immune deconvolution across more than 6,000 CNS tumors has demonstrated meaningful variation in TIME composition across tumor types, subtypes, grades, and anatomical locations, with several immune signatures associated with survival outcomes^12^. In ependymoma, crosstalk analyses integrating single-cell RNA sequencing data from tens of thousands of individual cells and validated across large patient cohorts have revealed elevated cell–cell interactions between malignant populations and tumor-infiltrating non-malignant cells, including interactions between neural stem cell-like tumor cells and microglia-derived populations, pointing to a functionally significant, immune compartment^13^. Recently, the application of spatial transcriptomics to PFA ependymoma has begun to map these cellular interactions onto defined tumor architectural regions, revealing the spatial organization of neoplastic subpopulations and their relationship with tumor progression^14^. However, despite these advances, the spatial distribution, functional state, and therapeutic relevance of rare lymphocyte infiltrates in pediatric ependymoma remain poorly characterized.

Here, we present a spatially resolved immune analysis of pediatric ependymoma using published Visium spatial transcriptomics data re-analyzed through an immune-focused computational framework. By integrating reference-guided deconvolution, spatial hotspot detection, functional lymphocyte state scoring, and ligand–receptor inference, we identified discrete lymphocyte-enriched niches in a subset of ependymoma sections, despite the overall low immune abundance. These niches are not randomly distributed but are preferentially associated with myeloid-rich and mesenchymal-like tumor regions, suggesting a model of spatially restricted immune access shaped by the local stromal and inflammatory context. Building on these findings, we extended the analysis to evaluate the spatial landscape of αβ and γδ T-cell immunotherapy opportunity, revealing heterogeneous but tractable therapeutic signatures across tumor sections. Together, this study provides a biological framework for immune restriction in pediatric ependymoma and a generalizable computational strategy for interrogating immune architecture and T-cell therapy susceptibility in low-infiltrated pediatric brain tumors.

## Results

### Immune landscape of pediatric CNS tumors reveals grade-dependent infiltration and location-specific lymphoid variation in ependymoma

To systematically characterize the immune microenvironment across pediatric CNS tumors, we applied LM7-based CIBERSORTx deconvolution to the OpenPBTA dataset, an open collaborative resource comprising RNA sequencing data from 1,074 pediatric brain tumors across 58 histological subtypes from 943 patients^15^. CIBERSORTx was selected for its robust performance in resolving immune cell composition from bulk gene expression data, including solid tumor tissues^16^. Rather than the standard LM22 signature matrix, we employed the optimized LM7 matrix, which was specifically developed to improve the resolution between closely related cytotoxic populations, namely CD8+ T cells, γδ T cells, and NK cells, a distinction underrepresented in conventional immune deconvolution frameworks^17^. Tumor types represented by fewer than three samples were excluded from the downstream analyses.

Ranking tumor types by cumulative inferred immune abundance revealed a broad spectrum of immune infiltration across a 19-tumor-type pediatric CNS landscape (Fig. 1a). Consistent with the well-established predominance of myeloid populations in CNS tumors, the monocyte/macrophage/dendritic cell (MoMaDC) subset comprised most of the inferred immune content across all histologies. Among all tumor types, desmoplastic infantile astrocytoma/ganglioglioma (DIA/DIG), meningioma (MNG), and pleomorphic xanthoastrocytoma (PXA), all tumors with low or absent malignant potential showed the highest cumulative immune abundance. Conversely, pineoblastoma (PB), oligodendroglioma (ODG), and embryonal tumors with multilayered rosettes (ETMR) displayed the lowest immune content across all subsets.

**Fig. 1:**
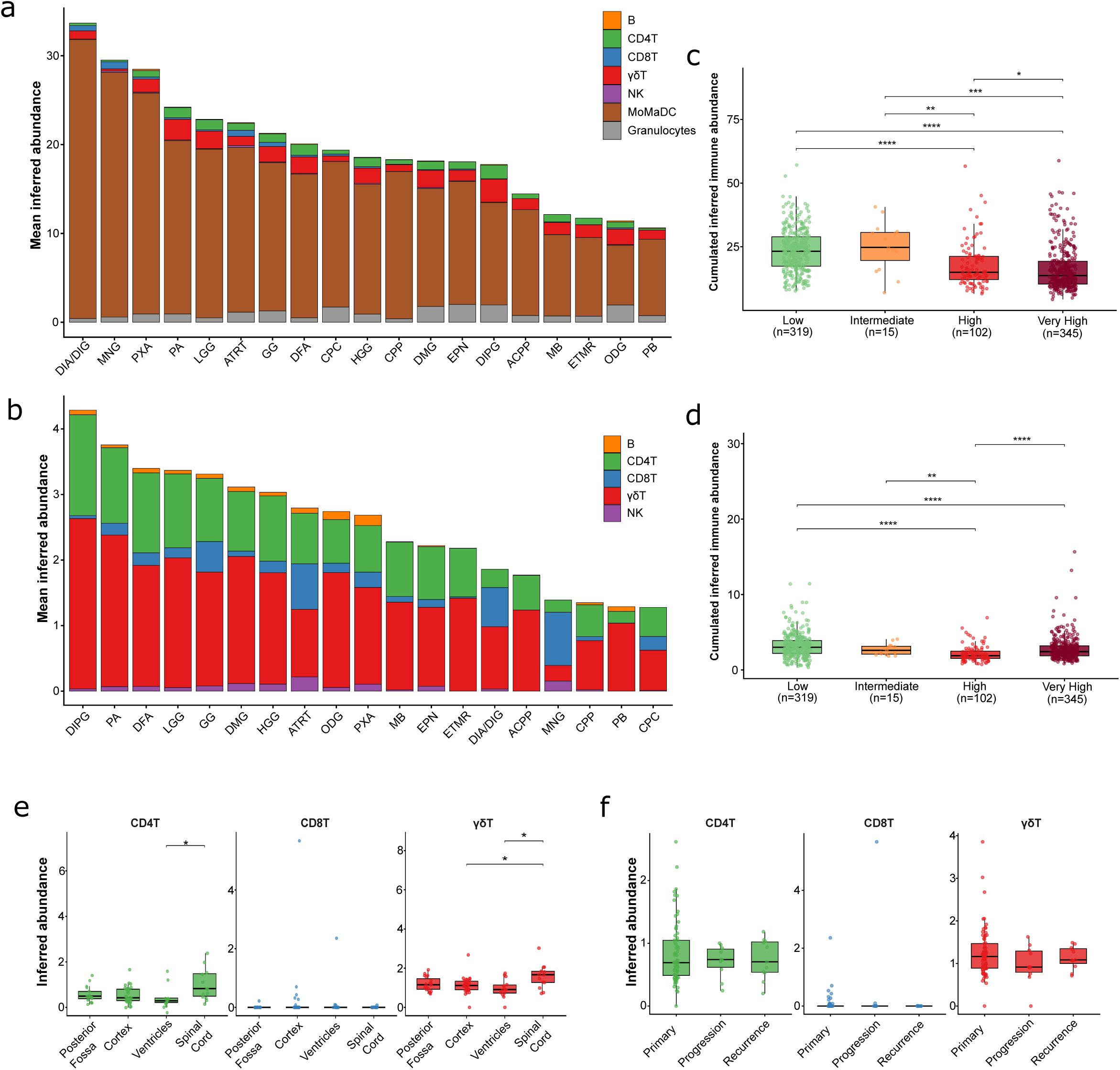
Pan-pediatric CNS tumor immune landscape reveals grade-dependent infiltration and location-specific lymphoid variation in ependymoma. **(a)** Mean inferred abundance of seven immune cell subsets: B cells, CD4+ T cells (CD4T), CD8+ T cells (CD8T), γδ T cells (γδT), NK cells, monocyte/macrophage/dendritic cells (MoMaDC), and granulocytes estimated by LM7-based CIBERSORTx deconvolution applied to bulk RNA sequencing data from the OpenPBTA dataset. Tumor types are ranked from highest to lowest total immune abundance. Only tumor types with n ≥ 3 samples are shown. Abbreviations: DIA/DIG, desmoplastic infantile astrocytoma/ganglioglioma; MNG, meningioma; PXA, pleomorphic xanthoastrocytoma; PA, pilocytic astrocytoma; LGG, low-grade glioma; ATRT, atypical teratoid/rhabdoid tumor; GG, ganglioglioma; DFA, diffuse fibrillary astrocytoma; CPC, choroid plexus carcinoma; HGG, high-grade glioma; CPP, choroid plexus papilloma; DMG, diffuse midline glioma; EPN, ependymoma; DIPG, diffuse intrinsic pontine glioma; ACPP, atypical choroid plexus papilloma; MB, medulloblastoma; ETMR, embryonal tumor with multilayered rosettes; ODG, oligodendroglioma; PB, pineoblastoma. **(b)** Mean inferred abundance of lymphoid subsets only (B, CD4T, CD8T, γδT, NK), re-ranked independently by total lymphoid abundance. The same tumor type abbreviations as in (a) apply. **(c)** Cumulated total inferred immune abundance (all seven subsets) stratified by tumor clinical aggressiveness category: Low (n=319), Intermediate (n=15), High (n=102), and Very High (n=345). Boxes indicate interquartile range (IQR), horizontal lines indicate median, and whiskers extend to 1.5× IQR. Individual samples are overlaid as jittered points. Statistical comparisons were performed using Kruskal–Wallis tests followed by pairwise Wilcoxon rank-sum tests with Benjamini–Hochberg correction. **(d)** Cumulated inferred lymphoid abundance (B, CD4T, CD8T, γδT, NK) stratified by clinical aggressiveness category, as in (c). Statistical testing as in (c). **(e)** Inferred abundance of CD4+ T cells, CD8+ T cells, and γδ T cells in ependymoma samples stratified by anatomical location: posterior fossa, cortex, ventricles, and spinal cord. Spinal cord ependymomas show significantly elevated CD4+ and γδ T cell abundance relative to intracranial locations. Statistical comparisons by Kruskal-Wallis test with Dunn’s post-hoc correction; * p<0.05. **(f)** Inferred abundance of CD4+ T cells, CD8+ T cells, and γδ T cells in ependymoma samples stratified by disease status: primary, progression, and recurrence. No statistically significant differences were observed across disease status categories.

Restricting the analysis to lymphoid subsets provided additional resolution of the adaptive and innate-like immune compartments (Fig. 1b). Across most pediatric CNS tumor types, γδ T cells constituted the largest lymphoid fraction, followed by CD4+ T cells. Meningioma was a notable exception, displaying a CD8+ T-cell-dominant lymphoid composition, consistent with previously reported histological and transcriptomic data for this tumor type^18,19^. Under this lymphoid-focused ranking, diffuse intrinsic pontine glioma (DIPG), pilocytic astrocytoma (PA), and diffuse fibrillary astrocytoma (DFA) emerged as the most lymphoid-infiltrated tumor types, whereas choroid plexus carcinoma (CPC), pineoblastoma (PB), and choroid plexus papilloma (CPP) showed the lowest lymphoid abundance.

Stratifying tumors by clinical aggressiveness demonstrated a significant inverse relationship between malignant potential and immune infiltration across the cohort. Both total immune abundance (Fig. 1c, Suppl. Table S1) and lymphoid abundance specifically (Fig. 1d) were significantly higher in low-aggressiveness tumors compared to high- and very high-aggressiveness categories (BH-adjusted p < 0.01 to p < 0.0001), suggesting that the immune-cold phenotype disproportionately characterizes aggressive pediatric CNS malignancies. A notable exception to this trend is DIPG, which ranked highest in inferred lymphoid abundance (Fig. 1b) despite being the most clinically aggressive tumor type in the cohort (WHO grade 4, median OS ∼11 months). This is likely attributable to the unique anatomical context of DIPG, a diffusely infiltrating brainstem tumor in close proximity to the meningeal compartment, which harbors resident T cell populations that may be captured in bulk transcriptomic data independently of genuine tumor infiltration^20^.

Having established these pan-tumor immune abundance patterns, we next focused on ependymoma, which ranked among the least immune-infiltrated tumor types in both analyses and carries a poor prognosis with no effective salvage therapy at relapse, to examine whether immune composition varied by anatomical location or disease status within this histological subtype. We stratified ependymoma samples by anatomical location (posterior fossa, cortex, ventricles, and spinal cord) and disease status (primary, progression, and recurrence) and independently assessed inferred CD4+, CD8+, and γδ T cell abundances. No significant difference in lymphoid infiltration was detected between the supratentorial compartments (cortex and ventricles) and the posterior fossa (Fig. 1e). However, consistent with the recognized biological and molecular distinctness of spinal ependymoma^21^, spinal cord ependymomas showed significantly elevated CD4+ and γδ T-cell abundance relative to intracranial ependymomas (Fig. 1e). No statistically significant difference in lymphoid infiltration was observed across disease status (primary, progression, or recurrence), suggesting that the immune composition in ependymoma is not substantially remodeled over the course of the disease (Fig. 1f).

Taken together, these data demonstrate that immune cell abundance in pediatric CNS tumors is primarily determined by tumor type and clinical aggressiveness, with more aggressive histology showing markedly reduced immune infiltration. Among ependymomas, only spinal cord localization was associated with a distinct lymphoid composition, characterized by elevated CD4+ and γδT cell abundance, a finding that may not be unrelated to the significantly better prognosis of spinal compared to intracranial ependymoma, where the immune-cold PFA subtype predominates. These findings motivated a more spatially resolved interrogation of the rare lymphoid infiltrates present in intracranial ependymoma, the subtype with the greatest clinical need.

### Spatial deconvolution identifies discrete lymphocyte-enriched niches in a subset of PFA ependymoma sections

To spatially resolve rare lymphocyte infiltrates in pediatric ependymoma, we applied cell-type deconvolution to published Visium spatial transcriptomic data from 14 PFA ependymoma sections (GSE195661)^14^. As a reference, we used an overlapping PFA ependymoma scRNA-seq cohort (GSE125969), re-annotated with simplified cell-type labels for downstream mapping^22^. Re-visualization of the scRNA-seq data by UMAP showed a discrete lymphocyte population that represented a markedly smaller fraction of cells than the myeloid compartment (Fig. 2a). Non-negative least-squares (NNLS) deconvolution was then performed using Semla with the complete set of cell-type groups present in the reduced reference, and the resulting Lymphocyte NNLS coefficient was carried forward for spatial analysis^23^.

**Fig 2:**
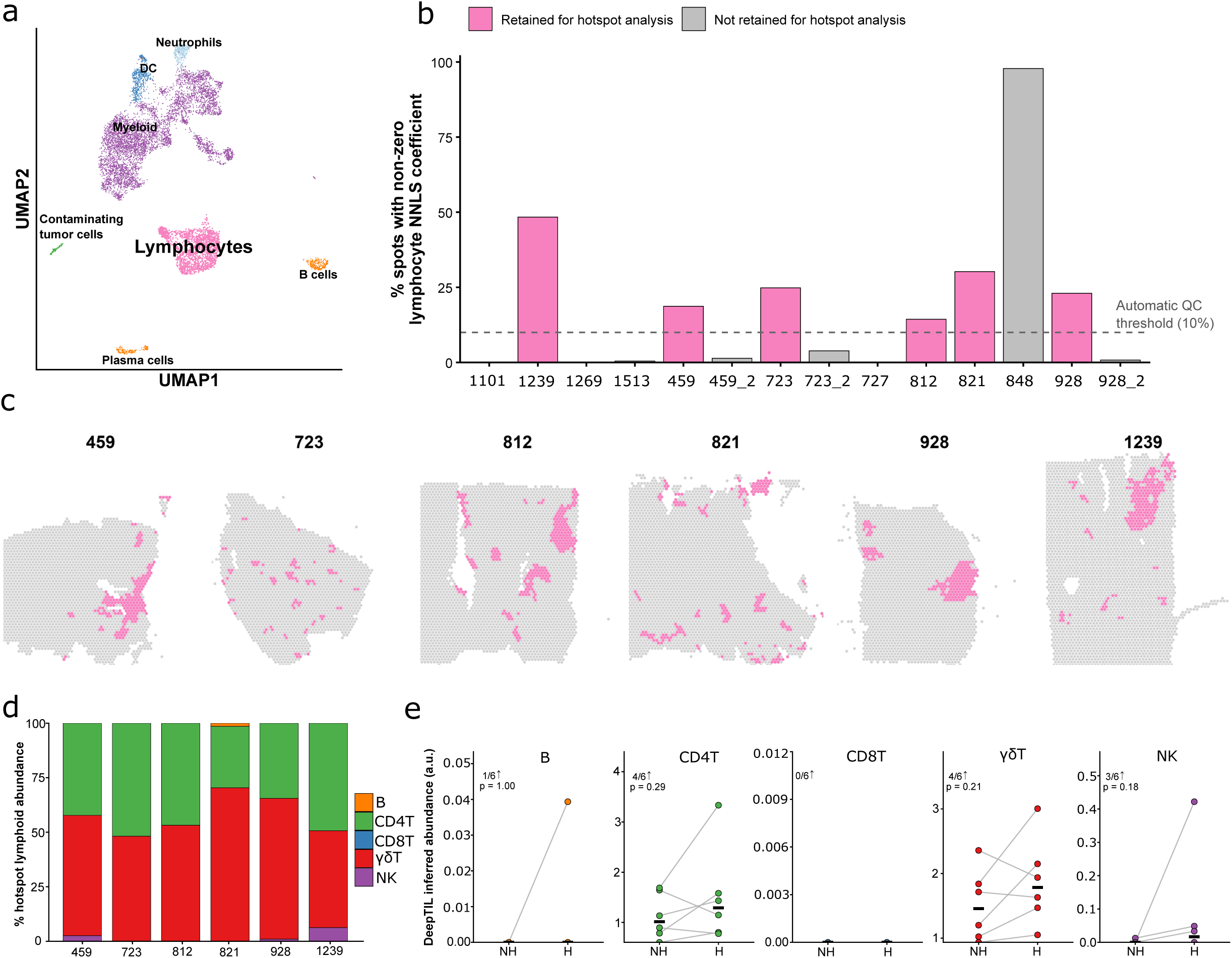
Spatial deconvolution identifies discrete lymphocyte-enriched niches in a subset of pediatric PFA ependymoma sections and reveals inter-patient heterogeneity in hotspot immune composition. **(a)** UMAP visualization of the single-cell RNA-seq reference dataset (GSE125969) used for spatial deconvolution, with cells colored by simplified cluster annotation: Myeloid, DC, Neutrophil, B-cells, Plasma-B-cells, contaminating Tumor cells and Lymphocytes. **(b)** Bar chart showing the percentage of Visium spots with a non-zero lymphocyte NNLS coefficient across all 14 spatial transcriptomic sections. Pink bars indicate the six sections retained for downstream hotspot analyses, while grey bars indicate the remaining sections. The dashed line denotes the 10% non-zero-spot lower-bound criterion used during automated section quality control. **(c)** Spatial maps of the six lymphocyte hotspot-positive sections, with hotspot spots highlighted in pink against the grey background of all in-tissue spots. **(d)** Stacked bar chart showing the lymphoid composition of hotspot pseudobulks inferred from LM7-based CIBERSORTx/DeepTIL deconvolution across the six hotspot-positive sections (459, 723, 812, 821, 928, 1239). For each section, inferred lymphoid abundance was normalized within the hotspot lymphoid compartment and displayed as the percentage contribution of five lymphoid populations: B cells, CD4 T cells, CD8 T cells, γδ T cells, and NK cells. **(e)** Paired plot showing LM7/DeepTIL-inferred abundance of the same five lymphoid populations in hotspot versus non-hotspot pseudobulks for each of the six hotspot-positive sections. Each point represents one section and grey lines connect matched hotspot and non-hotspot pseudobulks from the same section. Black horizontal bars indicate the median across sections. The annotation above each panel indicates the number of sections showing higher abundance in hotspots (x/6 ↑) together with the paired Wilcoxon signed-rank test p-value. Note that y-axis scales differ between panels.

Across the 14 spatial sections, the proportion of spots with a non-zero Lymphocyte NNLS coefficient varied substantially (Fig. 2b). Seven sections failed the predefined automated QC criteria, while section 848 passed the numerical screen but showed near-uniform lymphocyte signal across the capture area rather than a spatially focal pattern and was therefore excluded after spatial inspection. Six sections (459, 723, 812, 821, 928 and 1239) were retained for downstream hotspot analysis. Within these six sections, lymphocyte-enriched hotspots were defined as spots simultaneously exceeding the section-specific 90th percentile of the Lymphocyte NNLS coefficient and the 90th percentile of its six-nearest-neighbor local mean. Hotspots formed spatially discrete foci rather than diffuse distributions across the tissue (Fig. 2c), supporting a model in which lymphocyte infiltration is locally concentrated within selected regions of PFA ependymoma.

To assess whether the limited representation of neoplastic states in the original immune-focused reference biased hotspot detection, we repeated NNLS deconvolution using a PFA-specific tumor-augmented reference. The spatial Lymphocyte-score pattern was largely preserved. Per-section correlations between the original and tumor-augmented Lymphocyte scores ranged from ρ = 0.66–0.96 for raw scores and ρ = 0.75–0.98 after six-nearest-neighbour spatial smoothing. Hotspot overlap was substantial in five of six sections, with Jaccard indices of 0.63–0.80 and 73.8–86.7% of the original hotspots recovered. Section 723 showed lower concordance (Jaccard = 0.36; 49.4% recovery). Together, these analyses indicate that the recurrent spatial hotspot pattern was generally robust to inclusion of a substantially expanded PFA-specific neoplastic reference.

To characterize the immune composition of lymphocyte hotspot regions, raw Visium counts from hotspot, and non-hotspot spots were aggregated separately within each section and analyzed using LM7-based CIBERSORTx deconvolution followed by SES/DeepTIL abundance estimation. Within hotspot pseudobulks, the inferred lymphoid compartment was dominated by CD4 T cell- and γδ T cell-associated signals (Fig. 2d). γδ T cell-associated abundance represented the largest fraction of the modeled lymphoid compartment in four of six sections (459, 812, 821, and 928), accounting for approximately 53–70% of lymphoid abundance in these samples, whereas CD4 T cell-associated signal predominated in sections 723 and 1239. B cell-and CD8 T cell-associated contributions were negligible, while NK-associated abundance remained minor, reaching its highest relative contribution in section 1239. A comparable γδ T cell-associated contribution was observed in an independent bulk RNA-seq cohort spanning all ependymoma subtypes analyzed with the same LM7 matrix (Fig. 1b).

Comparison of paired hotspot and non-hotspot pseudobulks showed relatively modest and heterogeneous changes in inferred lymphoid abundance (Fig. 2e). CD4 T cell- and γδ T cell-associated abundance was higher in hotspots in four of six sections for each population (paired Wilcoxon, p = 0.29 and p = 0.21, respectively), whereas NK-associated abundance increased in three of six sections (p = 0.18). Together, these results indicate that the spatial hotspots represent locally slightly elevated lymphocyte-associated regions within an otherwise sparsely infiltrated tumor environment.

### Lymphocyte-enriched hotspots preferentially associate with myeloid and mesenchymal tumor zones and are depleted from epithelial neighborhoods

We first annotated each Visium spot with tumor region identity. Using the top marker genes of each cell cluster reported in the original Visium dataset publication, we applied module scoring to define four spatial zone programs per section: Epithelial, Mesenchymal, Vascular, and Myeloid; spots without a sufficiently strong assignment to any of these programs were classified as Uncertain (Fig. 3a, Suppl. Table S2a/b)^14^. This zone annotation framework was applied to all six hotspot-positive sections (459, 723, 812, 821, 928, and 1239), allowing for a systematic comparison of hotspot spatial positioning relative to the tumor architecture.

**Fig 3:**
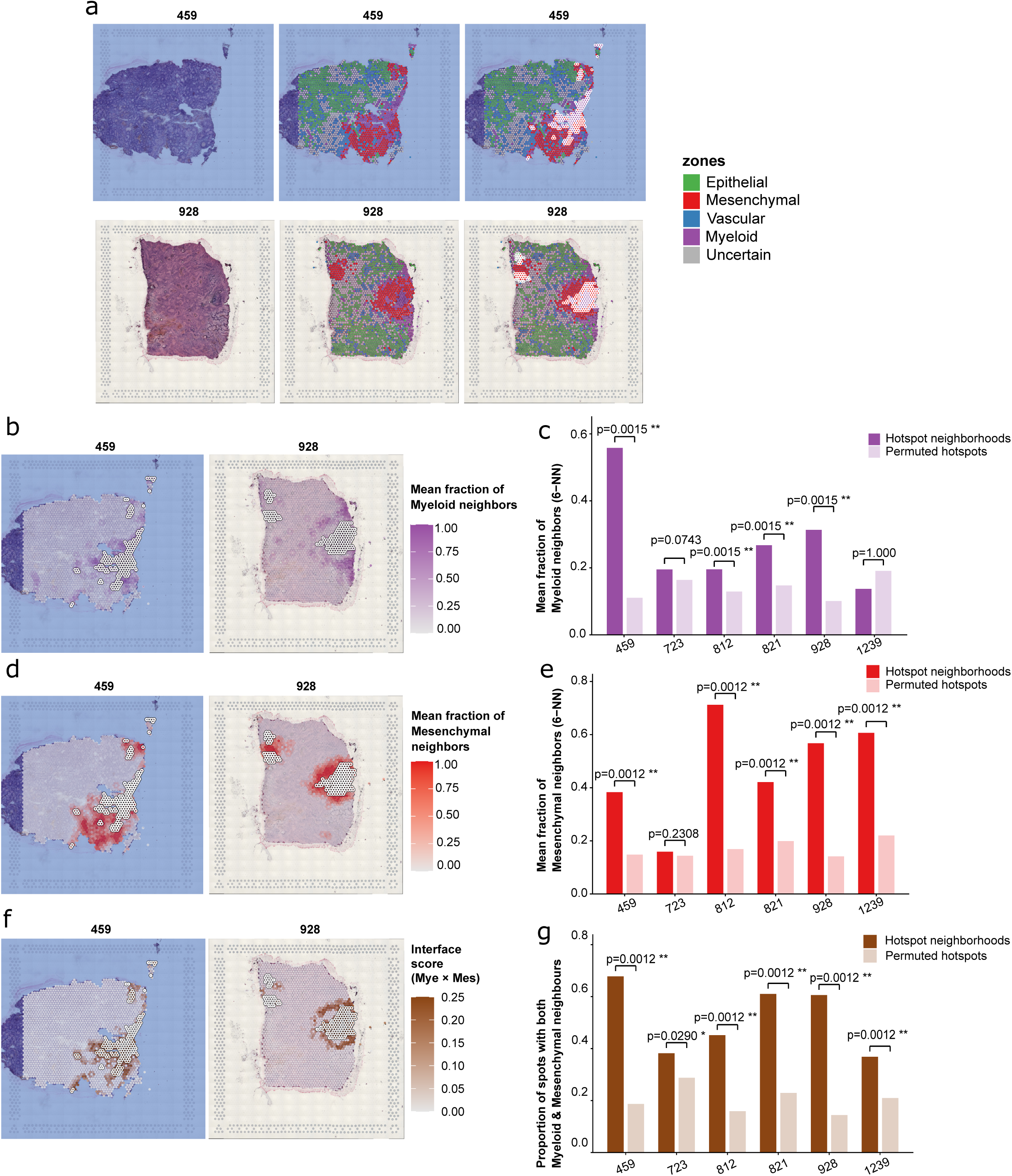
Lymphocyte-enriched hotspots are spatially confined to myeloid and mesenchymal tumor zones and preferentially localize at their shared interface. **(a)** Representative spatial maps of sections 459 (top row) and 928 (bottom row), each shown as three panels from left to right: HCE histology, tumor zone annotation, and zone map with lymphocyte hotspot spots overlaid. Zones are color-coded as follows: epithelial (green), mesenchymal (red), vascular (blue), myeloid (purple), uncertain (grey). Hotspot spots are displayed as white ring overlays. **(b)** Spatial maps of mean fraction of myeloid neighbors (6-nearest-neighbour radius) per spot for sections 459 and 928, illustrating enrichment of myeloid context adjacent to lymphocyte hotspot areas. **(c)** Bar chart comparing the mean fraction of myeloid neighbors in observed hotspot spots (dark purple) versus 1,000 permutations of hotspot labels (light purple) for each of the six hotspot-positive sections. Myeloid enrichment is significant in four out of six sections after multiple-testing correction. Significance was assessed using empirical permutation *P* values followed by Benjamini–Hochberg correction; ** BH-adjusted *P* < 0.005. **(d)** Spatial maps of mean fraction of Mesenchymal neighbors per spot for sections 459 and 928, illustrating co-localization of mesenchymal programs with lymphocyte hotspot areas. **(e)** Bar chart comparing the observed versus permuted mean fraction of mesenchymal neighbors across the six hotspot-positive sections, as in (c). Mesenchymal enrichment is significant in five out of six sections after multiple-testing correction. Significance was assessed using empirical permutation *P* values followed by Benjamini–Hochberg correction; ** BH-adjusted *P* < 0.005. **(f)** Spatial maps of the myeloid–mesenchymal interface score (product of myeloid neighbor fraction × mesenchymal neighbor fraction) per spot for sections 459 and 928. Higher scores indicate spots simultaneously embedded in both myeloid and mesenchymal neighborhoods. Hotspot spots (white rings) concentrate at regions of highest interface score. **(g)** Bar chart comparing the proportion of hotspot spots simultaneously adjacent to both myeloid and mesenchymal neighbors (observed, dark brown) versus 1,000 permutations of hotspot labels (light brown) across the six hotspot-positive sections. The co-neighborhood interface is significantly enriched in hotspot spots in all six sections after multiple-testing correction, with observed proportions consistently 2–4-fold above the permuted null. Significance was assessed using empirical permutation *P* values followed by Benjamini–Hochberg correction; * BH-adjusted *P* < 0.05, ** BH-adjusted *P* < 0.005.

We next assessed the local spatial context of lymphocyte-enriched hotspots using permutation-based neighborhood analysis. For each section, we quantified the mean fraction of zone-specific neighbors among the six nearest spatial neighbors of each hotspot spot and compared the observed hotspot-associated neighborhood composition with a null distribution generated from 1,000 random permutations of hotspot labels within the same section, preserving the number of hotspot spots. This section-specific analysis tested whether hotspots were preferentially positioned adjacent to Myeloid, Mesenchymal, Vascular, or Epithelial regions beyond that expected from the underlying spatial distribution of each zone. Empirical permutation *P* values were corrected across the six sections using the Benjamini–Hochberg procedure separately for each spatial test.

Myeloid neighbor enrichment in hotspot neighborhoods was significant in four out of six sections (459, 812, 821, and 928: all adjusted p = 0.0015; Fig. 3b/c), with section 723 showing a non-significant trend (adjusted p = 0.0743) and section 1239 showing no enrichment (adjusted p = 1.000). Mesenchymal neighbor enrichment was the most consistent finding, reaching significance in five out of six sections (459, 812, 821, 928, and 1239: all adjusted p = 0.0012; Fig. 3d/e), with only section 723 falling below significance (adjusted p = 0.2308). To further test whether hotspot spots specifically occupy the interface between myeloid and mesenchymal programs simultaneously rather than simply being near either zone independently, we computed a co-neighborhood interface score for each spot, defined as the product of its myeloid and mesenchymal neighbor fractions. The proportion of hotspot spots at this dual interface was significantly higher than permuted expectation across all six sections (five at adjusted p = 0.0012, section 723 at adjusted p = 0.029; Fig. 3f/g), with observed proportions consistently 2–4-fold above the null. This demonstrates that lymphocyte hotspots are not merely adjacent to myeloid or mesenchymal zones individually but are preferentially embedded at their shared spatial boundaries.

Complementary analyses confirmed that hotspot spots were consistently depleted of Epithelial neighbors across all six sections (all adjusted p = 0.001; Fig. S1a/b) and showed no enrichment for Vascular neighbors in five out of six sections (Fig. S1c/d), arguing against a perivascular model of lymphocyte infiltration. Section 723 showed consistently weaker or absent myeloid and mesenchymal neighborhood enrichment across all analyses, with its hotspot spots failing to localize to a coherent myeloid–mesenchymal interface (Fig. S1e). The overall pattern of myeloid–mesenchymal co-localization with lymphocyte hotspots was otherwise reproducible across sections despite inter-patient heterogeneity in zone composition.

To assess whether the co-enrichment of hotspots in both the mesenchymal and myeloid zones reflects shared biological programs between these two compartments, we computed the gene set overlap between zone-defining marker signatures. The mesenchymal and myeloid signatures shared 91 genes (Jaccard index = 0.23; 29.7% of the mesenchymal signature; 50.0% of the myeloid signature), including key inflammatory mediators such as *SPP1*, *CCL2*, *CCL5*, *CXCL1/2/3/5/6/8*, *VEGFA*, *MMP7*, *ICAM1*, *VCAM1*, and *IL32* (Suppl. Table S2c/d). These programs are associated with inflammatory activation, hypoxia response, and extracellular matrix remodeling, consistent with both populations occupying the same tumor microenvironment niche. By contrast, the lymphocyte signature shared only 3 genes with the Mesenchymal signature (*CCL5*, *IFITM1*, *IL32*; Jaccard = 0.006) and 2 genes with the Myeloid signature (*CCL5*, *IL32*; Jaccard = 0.005), indicating that lymphocyte hotspot identity is not driven by substantial direct gene overlap with the zone programs used to define their spatial context. These overlapping inflammatory programs between the mesenchymal and myeloid compartments, combined with their shared spatial association with lymphocyte hotspots, are consistent with a model in which a myeloid–mesenchymal interface shapes immune access and lymphocyte positioning in PFA ependymomas.

### Lymphocyte hotspots display inflammatory and tissue-retention-associated states and are bordered by SPP1–integrin, collagen–integrin, and APP–CD74 ligand-receptor axes from myeloid and mesenchymal zones

To characterize the functional identity of lymphocytes within the detected spatial hotspots, we applied UCell scoring to five curated functional state programs: cytotoxic-associated (*NKG7*, *PRF1*, *GZMB*, *GNLY*, *CTSW*, *KLRD1*, *KLRK1*, *XCL1*, *XCL2*, *CCL5*, *IFNG*), inflammatory-associated (*IFNG*, *TNF*, *CCL4*, *CCL5*, *XCL1*, *XCL2*, *NFKBIA*, *IRF1*, *STAT1*), exhausted-associated (*PDCD1*, *LAG3*, *TIGIT*, *HAVCR2*, *CTLA4*, *TOX*, *ENTPD1*, *CXCL13*), IL17-like-associated (*RORC*, *CCR6*, *KLRB1*, *IL7R*, *IL17A*, *IL17F*, *AREG*), and retention-associated (*CD44*, *ITGAE*, *CXCR3*, *CXCRc*, *CDcS*, *CCL5*). The retention-associated program was designed to capture a tissue-residency/tissue-retention transcriptional phenotype rather than to establish physical residence, antigen experience, or absence of cytotoxic activity. Hotspots meeting the confidence criteria were assigned the eligible state with the highest percentile-normalized score; weakly supported or ambiguous hotspots were left unassigned (Fig. 4a/b).

**Fig. 4.**
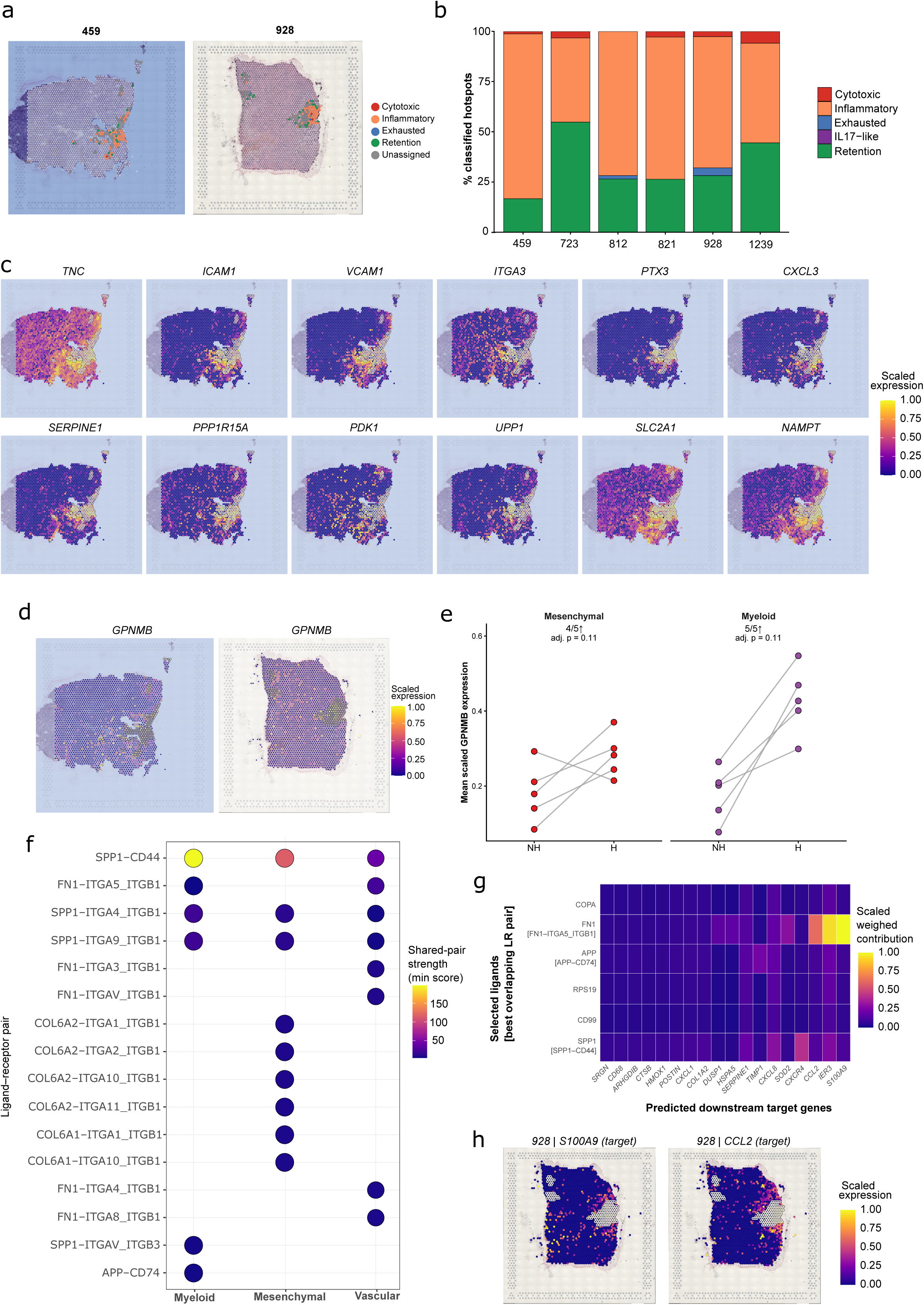
Lymphocyte hotspots display inflammatory and tissue-retention-associated states and are embedded within spatially organized ligand–receptor niches. **(a)** Representative spatial maps of sections 459 and 928 showing lymphocyte hotspot spots colored according to their confidently assigned dominant functional state. Functional states were scored using UCell signatures for cytotoxic, inflammatory, exhausted, IL17-like, and retention programs. A state was assigned only when the corresponding signature reached at least the 50th percentile of its section-specific hotspot distribution, at least two signature genes were detected in the individual spot, and the highest eligible state exceeded the second-ranked eligible state by at least0.05 on the percentile-normalized 0–1 scale. Hotspots not meeting these criteria were classified as unassigned and are shown in grey. Non-hotspot in-tissue spots are shown in light purple. (**b**) Relative distribution of confidently classified hotspot functional states across the six lymphocyte hotspot-positive sections (459, 723, 812, 821, 928, and 1239). Unassigned hotspots were excluded from the plotted proportions, and frequencies were renormalized to the total number of confidently classified hotspots within each section. The proportion of unassigned hotspots was retained separately for quality control. (**c**) Spatial expression maps of twelve representative niche-associated genes in section 459, overlaid on the HCE image, spanning adhesion and retention (*TNC, ICAM1, VCAM1, ITGA3*), inflammatory and chemoattractant (*PTX3, CXCL3*), and metabolic stress and hypoxia (*SERPINE1, PPP1R15A, PDK1, UPP1, SLC2A1, NAMPT*) programs. For visualization, expression of each gene was capped at its section-specific 99th percentile and scaled to this value. White-ringed spots indicate lymphocyte hotspots. (**d**) Spatial distribution of GPNMB expression in representative sections 459 and 928, overlaid on the corresponding HCE images. Expression was capped at the section-specific 99th percentile and scaled to this value, with values ranging from 0 to 1. White-ringed spots indicate lymphocyte hotspots. (**e**) Section-wise comparison of mean scaled GPNMB expression between hotspot and non-hotspot spots within Mesenchymal and Myeloid zones. Analyses were restricted to sections displaying the recurrent hotspot zone-confinement pattern and to section–zone combinations containing at least 10 hotspot and 10 non-hotspot spots. Each connected pair represents one section. GPNMB expression was higher in hotspot regions in 4/5 Mesenchymal sections and 5/5 Myeloid sections (paired Wilcoxon); neither comparison remained significant after Benjamini–Hochberg correction across the two zones (adjusted p = 0.11 for both). (**f**) Ligand–receptor (LR) interactions supported independently by both CellChatDB and CellPhoneDB, shown according to the spatial zone of the sender spot (Myeloid, Mesenchymal, or Vascular). Only pairs retained among the top-ranked interactions in both databases are displayed, providing a conservative set of interactions supported by two independent LR resources. Fill color represents the conservative shared interaction score, defined from the lower interaction score across CellChatDB and CellPhoneDB, whereas dot size represents the proportion of included sections supporting the interaction. LR pairs are shown on the y-axis and sender zone on the x-axis. (**g**) NicheNet analysis of candidate signaling at the Myeloid–hotspot interface. Candidate ligands were restricted to Myeloid LR interactions supported by both CellChatDB and CellPhoneDB and prioritized according to NicheNet ligand-activity AUC. Rows represent selected candidate ligands, annotated with their best-supported receptor, and columns represent predicted downstream target genes upregulated in hotspot spots neighboring the Myeloid zone. For each ligand–target pair, the weighted contribution was calculated as the NicheNet ligand–target regulatory potential multiplied by a receiver-gene weight defined as the mean log2 fold-change multiplied by the number of sections in which the target gene was upregulated. Values were scaled to the maximum weighted contribution across the displayed matrix for visualization. (**h**) Spatial expression of the NicheNet-predicted target genes S100A9 and CCL2 in section 928, shown as scaled expression over the corresponding tissue section.

After applying confidence criteria for functional-state assignment, 65.3–76.4% of hotspot spots per section remained unassigned, reflecting insufficient transcriptional support for a confident state call. Among confidently classified hotspots, an inflammatory-associated program was the predominant state in all six sections, accounting for 51.9–76.5% of classified spots (section 459: 76.5%; 723: 61.9%; 812: 68.4%; 821: 70.8%; 928: 61.0%; 1239: 51.9%). A retention-associated program represented the second most frequent assignment, ranging from 21.6% to 41.6% across sections. Cytotoxic and exhausted states were uncommon, together accounting for only a small fraction of confidently classified hotspots, while no hotspot was confidently assigned to the IL17-like state (Fig. 4a,b).

Thus, the confidence-filtered analysis identifies inflammatory- and retention-associated transcriptional programs as the principal states supported within lymphocyte hotspots, with comparatively limited evidence for cytotoxic or exhausted programs. The predominance of inflammatory-associated assignments together with the recurrent spatial localization of hotspots at myeloid–mesenchymal interfaces is consistent with a locally activated but non-classically cytotoxic immune environment, although the functional consequences of these transcriptional states cannot be inferred from spatial transcriptomic scoring alone.

To contextualize the functional states in relation to the surrounding tumor microenvironment, we visualized the spatial expression of representative genes associated with retention-permissive and metabolic stress programs across section 459, selected as the representative section given its large hotspot cluster (n = 180 spots) and spatially coherent hotspot architecture (Fig. 4c). These genes were chosen to characterize the microenvironmental context surrounding the hotspots rather than the hotspot lymphocytes themselves, and their expression therefore reflects the biology of the adjacent myeloid and mesenchymal zone cells rather than the functional state of the infiltrating lymphocytes. Adhesion and vascular cell adhesion molecules, including *TNC*, *ICAM1*, *VCAM1*, and *ITGA3*, showed enriched expression in the mesenchymal and myeloid-dominant regions of the tissue, co-localizing with the hotspot zone, consistent with a microenvironment that promotes lymphocyte adhesion and retention rather than transient trafficking. Inflammatory and chemoattractant markers including *PTX3* and *CXCL3* showed spatially concordant patterns in the same zone cells. Genes associated with metabolic stress and hypoxia adaptation (*SERPINE1*, *PPP1R15A*, *PDK1*, *UPP1*, *SLC2A1*, and *NAMPT*) displayed broad expression across the mesenchymal-enriched area, consistent with the hypoxia- and Warburg-associated programs identified in the shared Mesenchymal–Myeloid gene core (Suppl. Table S2c). Together, these expression patterns indicate that lymphocyte-enriched hotspots are embedded within a spatially coherent adhesive, inflammatory, and metabolically stressed niche where the inflammatory character derives from the surrounding myeloid and mesenchymal zone cells. We then examined the spatial expression of *GPNMB*, a gene recently identified as a dual-compartment antigen expressed in both immunosuppressive myeloid cells and mesenchymal-like tumor states in glioblastoma^24^. Spatial mapping of scaled *GPNMB* expression across sections 459 and 928 revealed striking co-localization with lymphocyte hotspot spots, particularly within myeloid- and mesenchymal-dominant zones (Fig. 4d). Section-wise comparison of *GPNMB* expression between hotspot and non-hotspot regions showed a consistent hotspot-associated increase within the Myeloid zone, with higher mean expression in all five retained sections (5/5; paired Wilcoxon, *p* = 0.059), and a similar but less consistent increase within the Mesenchymal zone (4/5 sections; *p* = 0.106). Neither comparison reached statistical significance after Benjamini–Hochberg correction (*adjusted p* = 0.106 for both) (Fig. 4e). These findings indicate that lymphocyte hotspots in PFA ependymoma are not only spatially confined to myeloid–mesenchymal interface niches but are also surrounded by cells expressing GPNMB, a marker associated with immunosuppressive myeloid reprogramming. This pattern is consistent with GPNMB marking an immunoregulatory myeloid–mesenchymal niche that may contribute to reduced functional capacity of infiltrating lymphocytes despite their spatial presence.

To systematically identify candidate intercellular signals from surrounding niche cells toward hotspot lymphocytes, we performed spatial ligand–receptor (LR) scoring independently using CellChatDB and CellPhoneDB. Detectable LR pairs from each resource were scored between Myeloid, Mesenchymal or Vascular sender spots and neighboring lymphocyte-hotspot receiver spots within a six-nearest-neighbor spatial graph. Five sections displaying the recurrent hotspot–zone confinement pattern (459, 812, 821, 928 and 1239) were included, with additional zone-specific exclusions where local zone representation was insufficient. LR scores represent spatially weighted transcriptomic interaction potential rather than confirmed signaling activity. The top 30 interactions per zone were retained independently for each database, and statistical robustness was assessed by hotspot-label permutation testing (Fig. S2a/b). Section 723 was excluded from this analysis because it did not display the recurrent neighborhood-confinement pattern observed in the other hotspot-positive sections (Fig. 3c/e/g).

The full top-30 rankings for each database independently are shown in Fig. S2c/d. We next focused on LR pairs that were independently retained among the top-ranked interactions in both CellChatDB and CellPhoneDB. Cross-zone comparison revealed distinct interaction landscapes (Fig. 4f). *SPP1–CD44* was the highest-scoring shared interaction in all three zones, with scores of 197.6 in Myeloid, 115.5 in Mesenchymal and 41.9 in Vascular regions, identifying a broadly recurrent candidate signaling axis. *FN1–ITGA5_ITGB1* was shared between the Myeloid and Vascular zones, whereas *FN1–ITGA4_ITGB1* and *FN1–ITGA8_ITGB1* were restricted to the Vascular cross-database consensus. In contrast, *COL6A2*–integrin interactions involving *ITGA1, ITGA2, ITGA10* and *ITGA11*, together with *COL6A1–ITGA1* and *COL6A1–ITGA10*, were restricted to the Mesenchymal consensus set, consistent with the prominent extracellular-matrix program of this compartment. Myeloid-specific consensus interactions included *APP–CD74* and *SPP1–ITGAV_ITGB3*.

To prioritize the most functionally relevant ligand-receptor interactions and link them to transcriptional changes in the hotspot receiver compartment, we applied NicheNet analysis with myeloid zone cells as senders and hotspot spots as receivers. Six ligands were retained after consensus filtering: *APP, FN1, SPP1, COPA, RPS1S, and CDSS* (Fig. 4g). For each ligand, NicheNet computed weighted regulatory potential scores linking the ligand to upregulated target genes in hotspot areas, yielding a scaled weighted contribution matrix. *APP,* whose highest-ranked overlapping LR pair was APP–CD74, showed the highest overall weighted regulatory contributions, with *S100AS* (NicheNet-predicted downstream target; weighted_scaled = 1.00, n = 2 sections, mean log2FC = 1.63) and *IER3* (weighted_scaled = 0.93, n = 2 sections, mean log2FC = 1.99) emerging as the top predicted downstream targets, followed by *CCL2* (weighted_scaled = 0.62, n = 1 section, mean log2FC = 1.95). S*PP1,* whose highest-ranked overlapping LR pair was SPP1–CD44, showed moderate contributions, with *CXCR4* as its primary predicted target (weighted_scaled = 0.41). *FN1* [via *FN1–ITGA5_ITGB1*] predicted *TIMP1* and *SERPINE1* as the top targets. COPA, RPS19, and CD99 exhibited weaker and less specific weighted contribution profiles. Spatial concordance in section 928, which had the highest *APP–CD74* LR score, confirmed the enriched expression of both *S100A9* and *CCL2* in the myeloid-dominant zone neighboring lymphocyte hotspots (Fig. 4h), providing spatial concordance between the *APP*-associated NicheNet target program and the *APP–CD74* candidate LR interaction.

Taken together, these data support a model in which PFA ependymoma lymphocyte hotspots are spatially associated with SPP1–CD44/integrin, collagen VI–integrin, and APP–CD74 signaling axes emanating from the adjacent myeloid and mesenchymal microenvironment, consistent with inflammatory- and tissue-retention-associated transcriptional states, with comparatively limited evidence for active cytotoxic engagement.

### Spatially resolved opportunity scoring reveals heterogeneous and compartmentalized αβ and γδ T cell recognition landscapes in ependymoma

To evaluate whether the ependymoma sections displayed transcriptional features compatible with αβ or γδ T cell-based immunotherapy, we developed a multi-component spatial opportunity scoring framework across all 14 Visium sections. Three gene programs were quantified at the spot level: an αβT ligand availability program capturing antigen-presentation machinery (HLA-A/B/C, B2M, TAP1/2, PSMB8/9, ERAP1/2, NLRC5, IRF1); a γδT ligand availability program capturing butyrophilin, phosphoantigen-pathway, stress-ligand, ephrin and adhesion features (BTN2A1, BTN3A1; MVK, PMVK, MVD, IDI1; MICA, MICB, ULBP2, ULBP3, RAET1E, RAET1G, RAET1L; EPHA2; PVR, NECTIN2, ICAM1); and a shared inhibitory program incorporating canonical immune checkpoints, immunosuppressive cytokines and mediators of the COX–PGE2, adenosine and vascular suppressive axes (CD274, PDCD1LG2, HLA-E, LGALS9, IDO1, TGFB1-3, IL10, VEGFA, NT5E, ENTPD1, PTGS2, PTGES, PTGER2/4). These programs measure the tumor-side ligand landscape and suppressive-program expression. Section-level net scores were calculated by subtracting the shared inhibitory score from the corresponding αβT or γδT ligand availability score, and sections were classified according to the relative balance between opportunity and inhibitory signals. Because gene expression was z-transformed across the pooled tumor spots of all sections, the resulting scores and categories are relative to this 14-section cohort rather than absolute. After applying the predefined 5% tumor-spot detection threshold, 12/12 αβT genes, 11/17 γδT genes, and 9/16 immunosuppressive genes were retained for scoring.

The resulting opportunity landscape showed substantial heterogeneity across the cohort (Fig. 5a). Sections 821, 1101, 723_2, 727 and 928 were classified as αβT-favorable; sections 848, 723, 1513 and 459 as γδT-favorable; sections 928_2 and 459_2 as Dual-opportunity; and sections 812 and 1269 as Immuno-cold. Section 1239 showed a markedly elevated inhibitory burden and was the only section classified as Suppressed (Fig. S3b). This reflected concurrent elevation across mechanisms rather than a single dominant axis: 1239 ranked highest in the cohort for COX-2, VEGF, HLA-E/NKG2A, adenosine and TGF-β, and second for galectin-9 (Fig. S3a). Several sections lay close to a category boundary, 928 and 727, within 0.02 of zero on the αβT axis, and the categories should therefore be read as a discretization of continuous scores rather than as discrete states.

**Fig. 5:**
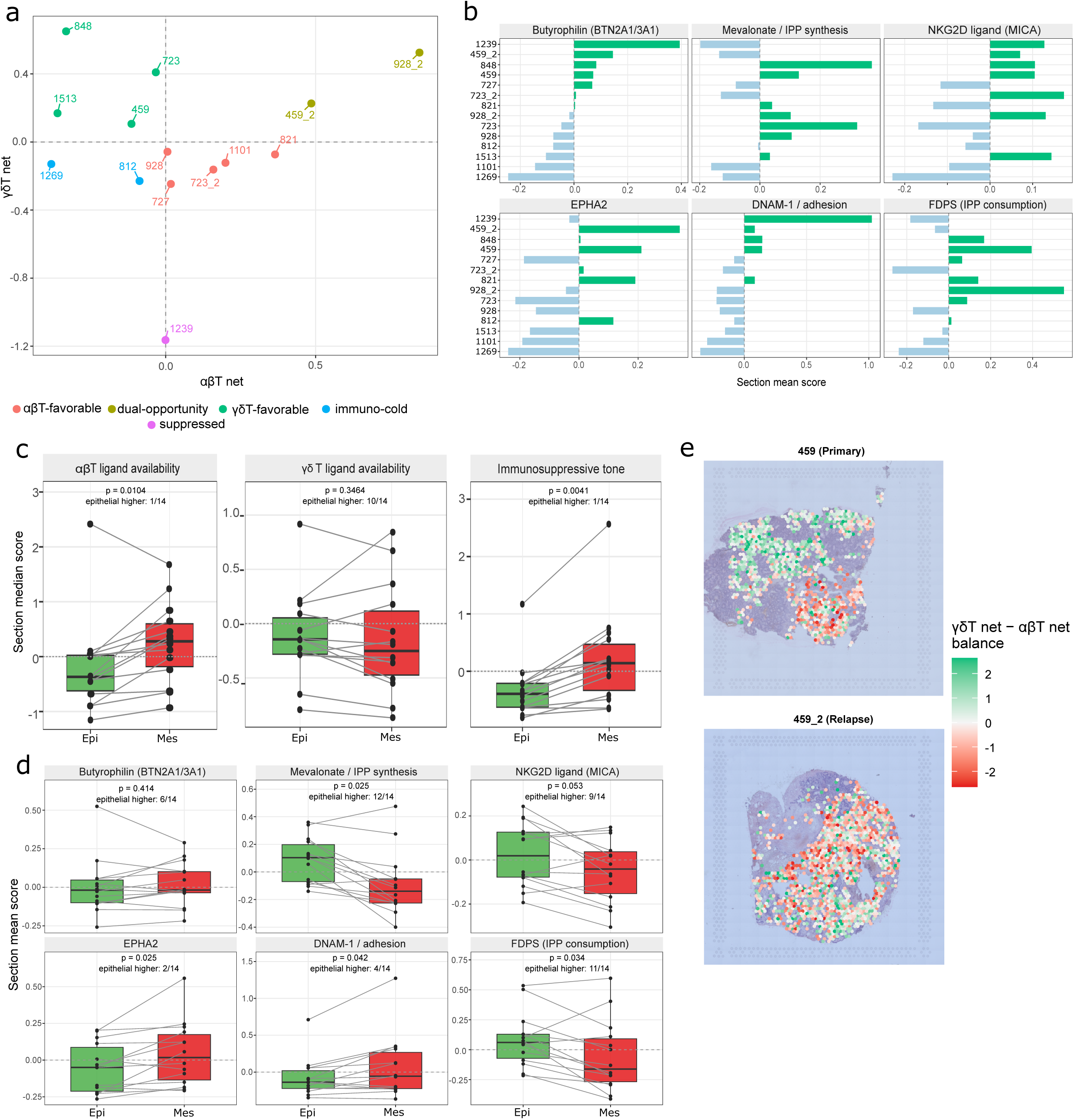
Spatial immunotherapy opportunity scoring reveals heterogeneous αβ and γδ T cell opportunity landscapes across and within PFA ependymoma sections. **(a)** αβ/γδT opportunity map across all 14 ependymoma sections. Each point represents one section, positioned by its median αβT_net score (x-axis) and γδT_net score (y-axis), derived as the corresponding core opportunity z-score minus the shared immunosuppressive program z-score. Sections are color-coded by opportunity category: αβT-favorable (positive αβT_net, negative γδT_net; red), γδT-favorable (negative αβT_net, positive γδT_net; green), Dual-opportunity (both positive; olive), Immuno-cold (both ≤0 with low immunosuppressive burden; cyan), and Suppressed (both ≤0 with high immunosuppressive burden; magenta). Scores are z-scaled within the cohort and therefore represent relative opportunity across the 14 sections. **(b)** Section-level γδT recognition modality scores across the same 14 ependymoma sections. Each bar represents the mean spot-level score for one section for six recognition-related modalities: Butyrophilin (BTN2A1/3A1), Mevalonate/IPP synthesis, NKG2D ligand (MICA), EPHA2, DNAM-1/adhesion, and FDPS (IPP consumption). Modalities were scored independently; the recognition modalities contributing to the γδT composite were equally weighted. FDPS was treated as a modifier and was not included in the γδT composite opportunity score. **(c)** Opportunity components stratified by tumor compartment (epithelial versus mesenchymal), computed per section and compared across the 14 sections (n = 14 pairs). Each point is one section’s median spot-level score within that compartment, and lines connect the two compartments of the same section. Epithelial and mesenchymal scores were compared by paired Wilcoxon signed-rank test; p values are Benjamini–Hochberg adjusted. **(d)** γδT recognition modalities stratified by tumor compartment (epithelial versus mesenchymal), computed per section and compared across the same 14 sections (n = 14 pairs). Each point is one section’s mean spot-level score within that compartment, and lines connect matched epithelial and mesenchymal values from the same section. Paired Wilcoxon signed-rank tests indicate the direction and significance of compartmental differences, with Benjamini–Hochberg-adjusted p values; y scales differ between panels, and no score is subtracted from another. FDPS (IPP consumption) is shown as a modifier and is not part of the γδT composite. **(e)** Spatial maps of the γδT-versus-αβT opportunity balance in primary section 459 (top) and its matched relapse section 459_2 (bottom) from the same patient. Each spot is colored according to the balance score, calculated as γδT_net − αβT_net. Positive values (green) indicate relative γδT opportunity, whereas negative values (red) indicate relative αβT opportunity. The immunosuppressive component cancels exactly in this contrast, allowing the maps to directly visualize the spatial balance between γδT and αβT recognition opportunity.

The three matched primary–relapse pairs each showed remodeling of the opportunity landscape at relapse: 459 shifted from γδT-favorable to Dual-opportunity, 928 from αβT-favorable to Dual-opportunity, and 723 from γδT-favorable to αβT-favorable. Decomposing the net scores into their components showed that this movement was driven from two directions (Fig. S3c): αβT ligand availability increased in all three patients, immunosuppressive tone decreased in two of three, and γδT ligand availability increased in only one. The consistency of the αβT increase is notable, but with three matched pairs no formal inference is possible, and the observation should be regarded as hypothesis-generating.

Recognition modalities differ between sections. The γδT program aggregates five recognition modalities, which showed distinct patterns of variation across the cohort (Fig. 5b). Mevalonate/IPP synthesis was highest in sections 848 and 723, butyrophilin and DNAM-1/adhesion in 1239, EPHA2 in 459_2, and FDPS in 928_2. No section scored uniformly high across modalities, indicating that a favorable composite score can arise through different recognition routes, a distinction with therapeutic consequences, since butyrophilin-dependent and NKG2D-dependent recognition are engaged by different strategies.

Within sections, we compared the three components between the epithelial and mesenchymal tumor compartments (Fig. 5c). αβT ligand availability was higher in the mesenchymal compartment in 13 of 14 sections (adjusted p = 0.010) and immunosuppressive tone in 13 of 14 (adjusted p = 0.004), whereas γδT ligand availability showed no compartment difference (adjusted p = 0.35).

αβT ligand availability and immunosuppressive tone followed the same mesenchymal gradient, indicating that these two features are spatially coupled in this tissue: the compartment with the highest antigen-presentation machinery also displayed the strongest immunosuppressive program. The individual suppressive mechanisms behaved consistently, with HLA-E/NKG2A, adenosine, TGF-β, COX-2 and VEGF, all higher in the mesenchymal compartment and only galectin-9 showing no difference (Fig. S3d). VEGFA and PTGS2 also contribute to the mesenchymal zone definition, so those two comparisons are not independent of the compartment assignment; the remaining mechanisms share no genes with it. Decomposing the γδT program into its modalities showed that the absence of a compartment difference reflects cancellation rather than uniformity (Fig. 5d). Mevalonate/IPP synthesis was higher in the epithelial compartment in 12 of 14 sections (adjusted p = 0.025) and NKG2D ligand in 9 of 14 (adjusted p = 0.053), whereas EPHA2 was higher in the mesenchymal compartment in 12 of 14 (adjusted p = 0.025) and DNAM-1/adhesion in 10 of 14 (adjusted p = 0.042); butyrophilin showed no compartment preference (adjusted p = 0.41). Because the composite weights each modality equally, these opposing gradients cancel. This has a direct implication: the compartment in which γδ T cells would encounter the strongest recognition signal depends on which modality is engaged. Phosphoantigen-dependent recognition would be favored in epithelial regions, whereas adhesion- and ephrin-dependent recognition would be favored in mesenchymal regions. One caveat qualifies the phosphoantigen interpretation, FDPS, which consumes IPP, was also higher in the epithelial compartment in 11 of 14 sections (adjusted p = 0.034), so elevated mevalonate-pathway expression does not by itself establish elevated phosphoantigen accumulation.

Mapping the balance between the two modalities per spot (γδT net − αβT net, in which the shared inhibitory term cancels exactly) showed that this balance is not uniform within a section (Fig. 5e). In the primary section 459, regions favoring γδ recognition occupied one contiguous portion of the tissue while αβ-favoring regions occupied another; in the matched relapse 459_2, the balance shifted toward αβ across most of the section, consistent with the section-level component change.

### Opportunity scores are embedded in distinct transcriptional contexts

To characterize the transcriptional programs co-varying with each opportunity score, genes were ranked by their association with the score within each section and the 14 rankings combined, with enrichment tested against Hallmark and Reactome gene sets. Because the scores are derived from expression, pathways containing the scoring genes themselves are expected to enrich; each cell is therefore annotated with whether the pathway shares genes with that score’s program, and with the number of sections in which the section-level enrichment agreed in sign with the combined result, cells below 75% concordance being drawn at reduced opacity.

The three composite scores occupied distinct transcriptional contexts (Fig. 6a). αβT ligand availability and immunosuppressive tone were associated with almost the same programs: interferon and antigen-presentation pathways, inflammatory and cytokine signaling, hypoxia, glycolysis and apoptosis, epithelial–mesenchymal transition and extracellular matrix organization; and both were negatively associated with oxidative ATP production. This near-complete overlap is consistent with the shared mesenchymal gradient described above and reinforces their spatial and transcriptional coupling. γδT ligand availability was associated with a largely non-overlapping set: cholesterol and lanosterol biosynthesis, dolichol-linked N-glycan precursor biosynthesis, and, among immune programs, only interferon-α response and TNF-α signaling. The isoprenoid enrichments contain the mevalonate genes used to build the score and are expected. The dolichol-linked N-glycan pathway is flagged as containing program genes, so it too cannot be read as independent discovery, although its enrichment is consistent with dolichol being synthesized from farnesyl pyrophosphate downstream of the same pathway.

**Fig 6:**
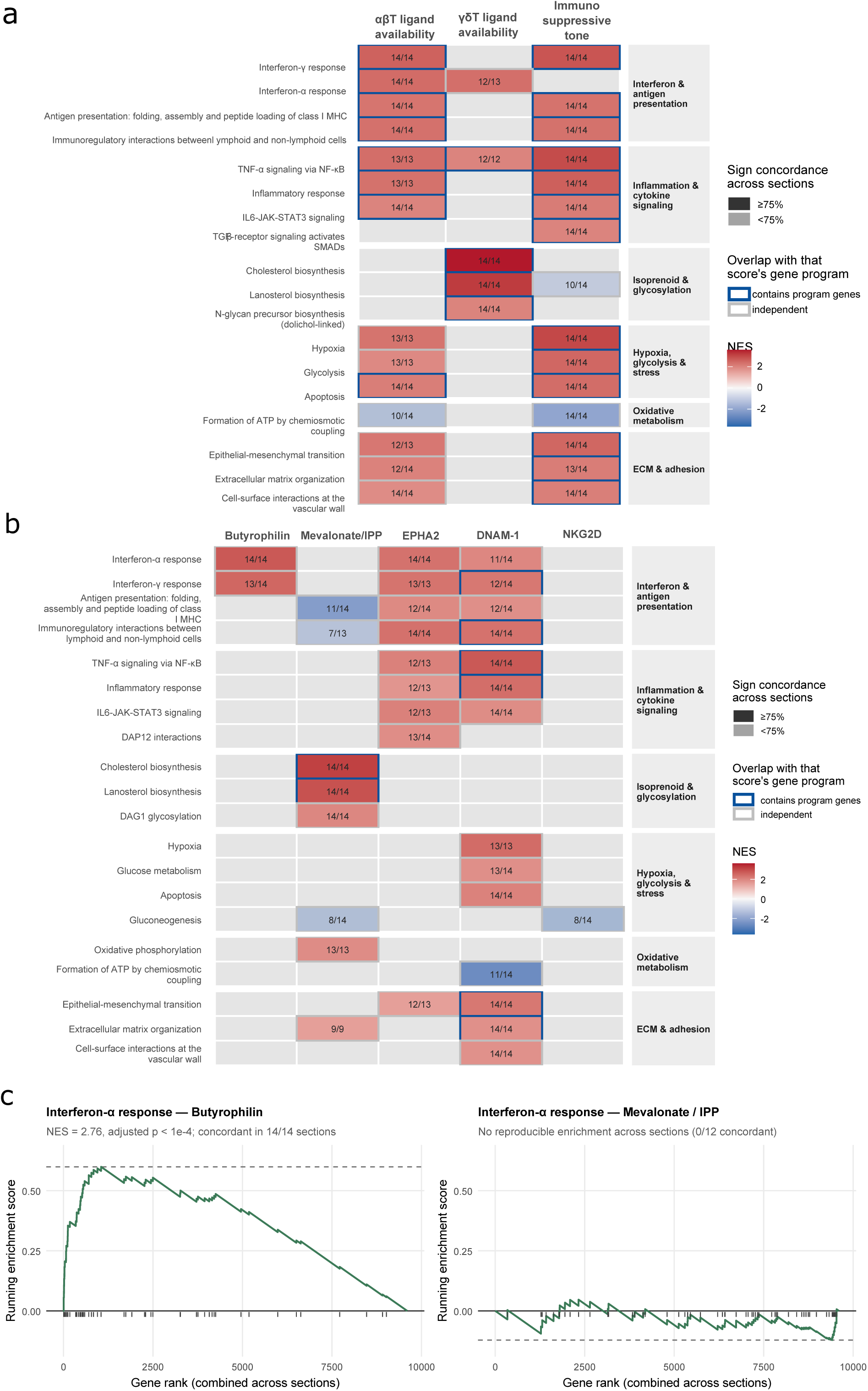
Pathway associations of T-cell opportunity scores and γδ T-cell recognition axes. **(a)** Gene set enrichment analysis (GSEA) of the three composite opportunity scores: αβ T-cell ligand availability, γδ T-cell ligand availability, and immunosuppressive tone. Selected biologically interpretable and non-redundant Hallmark and Reactome pathways among significant collapsePathways main pathways are shown. Tile color represents the normalized enrichment score (NES), with red and blue indicating positive and negative enrichment, respectively; grey cells indicate pathways not significantly enriched for the corresponding score. Numbers within significant cells indicate the number of sections showing the same direction of association over the number of sections tested. Reduced opacity denotes sign concordance <75% across sections. Tile borders indicate whether the pathway contains genes used to construct the corresponding opportunity score (blue) or is independent of the score gene program (grey). Pathways are grouped by biological function. **(b)** GSEA of the five γδ T-cell mechanism axes: butyrophilin, mevalonate/IPP, EPHA2, DNAM-1/adhesion, and NKG2D ligands. Encoding is as in (a). The heatmap highlights distinct pathway associations across γδ T-cell recognition mechanisms, including interferon-associated enrichment for the butyrophilin axis, sterol/isoprenoid metabolism for the mevalonate/IPP axis, and inflammatory, hypoxic, and extracellular-matrix programs associated with EPHA2 and DNAM-1/adhesion. The NKG2D-ligand axis showed little reproducible pathway-level enrichment, with gluconeogenesis representing the only significant main pathway retained and showing low cross-section sign concordance. **(c)** Running enrichment profiles for the Hallmark interferon-α response gene set along the combined gene rankings for the butyrophilin and mevalonate/IPP axes. The butyrophilin axis showed strong positive enrichment (NES = 2.76, adjusted *p* < 1 × 10⁻⁴), concordant in 14/14 sections, whereas no reproducible enrichment was observed for the mevalonate/IPP axis (0/12 sections concordant). Vertical ticks indicate positions of interferon-α response genes in the ranked gene list, and dashed horizontal lines indicate the maximal absolute running enrichment score. Both panels are shown on the same y-axis scale to enable direct comparison. For all heatmaps, significance was defined as Benjamini–Hochberg-adjusted *p* < 0.05. Only collapsePathways main pathways were considered for display.

Resolving the γδT program into its five modalities showed that the composite conceals two divergent contexts (Fig. 6b). Butyrophilin was associated only with interferon-α (14/14 sections) and interferon-γ (13/14) responses. Mevalonate/IPP showed the reciprocal pattern: cholesterol and lanosterol biosynthesis and DAG1 glycosylation, oxidative phosphorylation, and *negative* associations with antigen presentation and with immunoregulatory interactions between lymphoid and non-lymphoid cells. The negative association with antigen presentation showed 11/14 section-level sign concordance, whereas the association with immunoregulatory interactions was less consistent (7/13 sections); the latter should therefore be interpreted cautiously. EPHA2 and DNAM-1 were both associated with broad inflammatory, interferon and ECM programs, with DNAM-1 additionally associated with hypoxia, glucose metabolism and apoptosis; note that ICAM1, PVR and NECTIN2 are themselves members of several of these gene sets, and those cells are marked accordingly. NKG2D ligands (MICA) yielded a single association across the entire analysis, a weakly concordant negative enrichment of gluconeogenesis (8/14).

Because butyrophilin presentation and phosphoantigen supply are both required for Vγ9Vδ2 recognition, we compared their transcriptional contexts for the same gene set directly (Fig. 6c). Interferon-α response was enriched with the butyrophilin axis at NES = 2.76, adjusted p < 1 × 10⁻⁴, with concordant sign in 14 of 14 sections, whereas the same pathway showed no reproducible enrichment with mevalonate/IPP, the section-level results agreeing in sign with the combined result in 0 of 12 sections. The two prerequisites for phosphoantigen-mediated Vγ9Vδ2 recognition, the presenting butyrophilin molecules and the metabolic pathway generating their ligand, therefore sit in unrelated transcriptional environments in this tumor. Butyrophilin expression therefore tracks an interferon-associated transcriptional context, whereas mevalonate-pathway expression tracks lipid and glycosylation metabolism and shows an inverse association with antigen-presentation programs. Together with their different compartmental patterns, mevalonate/IPP being enriched in epithelial regions while butyrophilin showed no compartment preference, these findings indicate that the two requirements for phosphoantigen-mediated Vγ9Vδ2 recognition are not consistently coupled across PFA ependymoma.

## Discussion

Pediatric ependymoma, including PFA, is conventionally regarded as immunologically cold, with a low mutational burden and sparse lymphocyte infiltration, limiting enthusiasm for immune checkpoint-based approaches. That same biology, however, motivates T cell strategies that do not depend on neoantigen load. For αβ T cells, the constraint is antigenic: with few somatic mutations, the relevant targets are not private neoantigens but shared tumor-associated antigens presented on MHC class I, for which HLA ligandome atlases in ependymoma and medulloblastoma now provide a defined substrate, and which are addressable by peptide or mRNA vaccination and by adoptive transfer of TCR-engineered cells^25^. γδ T cells circumvent the constraint altogether: recognition through butyrophilin/phosphoantigen, NKG2D stress ligands, ephrin receptors and DNAM-1 ligands are MHC-unrestricted and reports metabolic and stress states of the tumor cell rather than its mutational history^26^.

Our spatially resolved analysis takes that view and asks where, within a tumor, each form of recognition is best supplied. Opportunity scoring across 14 sections showed that PFA ependymoma is not uniformly unsuitable for T cell-based approaches: individual sections displayed transcriptional features favoring αβ or γδ recognition, or both, with marked heterogeneity between and within tumors. That heterogeneity is compartmentalized rather than random; antigen-presentation machinery and immunosuppressive tone rise together in mesenchymal regions, while the modalities supporting γδ recognition are distributed in opposing directions across the same compartments. Where lymphocyte-associated signal was detectable at all, it was spatially restricted to myeloid- and mesenchymal-rich zones and largely absent from epithelial regions, and ligand–receptor analysis nominated a small set of candidate axes, SPP1–CD44, FN1–integrin, collagen VI–integrin and APP–CD74, that plausibly underlie that restriction. Together these findings reframe PFA ependymoma not as a monolithic immune desert, but as a tumor in which recognition potential and lymphocyte access are separately organized in space, a distinction with direct consequences for how immunotherapy should be designed and sequenced in this disease.

The ligand–receptor analysis identified four mechanistically distinct but spatially convergent candidate axes that may contribute to restricted lymphocyte access and function within the myeloid–mesenchymal niche, each suggesting a potential therapeutic vulnerability. SPP1–CD44 emerged as the highest-scoring pair across both myeloid and mesenchymal zones. Myeloid-derived SPP1 suppresses T cell proliferation, IFN-γ production, and early activation markers in a dose-dependent manner via CD44, and activates PI3K/AKT, MAPK/ERK, and FAK cascades that recruit Tregs and MDSCs; its blockade reverses immunosuppressive TME conditions and enhances CAR T cell responses in pediatric CNS tumors. SPP1–CD44 signaling has further been shown to mediate tumor–immune cell interactions leading to Immune checkpoint Inhibitor (ICI) resistance, providing a rationale for disrupting this axis therapeutically^27^. FN1–ITGA5_ITGB1 was detected in the Myeloid and Vascular zones, with additional FN1–integrin variants exclusive to the Vascular zone, pointing to a perivascular physical barrier rather than a molecular checkpoint. Targeting α5β1 integrin to disrupt mature fibronectin ECM reduces fibronectin fibril formation, enhances CD8+ T cell transendothelial migration, and improves PD-L1 blockade efficacy, positioning this axis upstream of and complementary to SPP1–CD44-mediated suppression^28^. COL6A1/COL6A2–integrin pairs were exclusively restricted to the mesenchymal zone, consistent with a structural entrapment model. Collagen VI forms a microfibrillar network at the basement membrane–interstitial matrix interface; collagen fragments in the cancer microenvironment mediate T cell suppression through LAIR-1, and multiple integrin pairs (α1β1, α2β1, α3β1, α10β1) link collagen VI ECM adhesion to regulation of T cell activation and TCR signaling, suggesting the dense mesenchymal matrix simultaneously immobilizes and suppresses lymphocytes at the hotspot boundary^29^. APP–CD74 was the only myeloid-exclusive axis, pointing to a dedicated macrophage-to-lymphocyte signaling event. The APP–CD74 axis drives M2-like TAM polarization in solid tumors, with functional assays confirming that APP and CD74 activation promote immunosuppressive macrophage programming^30^; in the CNS context specifically, increased APP is associated with broad suppression of adaptive and innate anti-tumor immunity including B cell-mediated and complement-dependent responses^31^. Its top NicheNet downstream targets: S100A9, IER3, and CCL2, converge on a coherent immunosuppressive program. S100A9 amplifies MDSC accumulation via TLR4/RAGE signaling; IER3 dampens NF-κB downregulation, locking the myeloid compartment in a non-resolving inflammatory state; and CCL2 creates a self-reinforcing myeloid recruitment loop through CCR2+ monocyte and MDSC chemotaxis^32,33^. A mechanism well-characterized in glioma, where CCR2 antagonism improves checkpoint efficacy, is supported by the spatial co-localization of CCL2 with the myeloid-adjacent zone in section 928 in ependymoma. These four axes are consistent with communication architectures identified by LR multi-omics in glioma, where myeloid-dominant immunosuppression and spatially organized immune exclusion are governed by reproducible TAM–tumor interaction programs^34^. Our ependymoma data extend this framework to a pediatric, immune-cold CNS tumor context not previously addressed by spatial LR approaches and suggest that unlocking the rare but spatially defined lymphocyte infiltrates identified here may benefit from combinatorial strategies targeting the ECM-associated, myeloid-polarization and chemokine-recruitment programs identified here rather than any single axis in isolation.

The opportunity categories identified here carry direct translational implications in the context of emerging immunotherapy data for pediatric CNS tumors. Four of the fourteen sections were classified as γδT-favorable, reflecting relative enrichment in butyrophilin ligands, NKG2D stress ligand (MICA), EPHA2, DNAM-1 ligands and the mevalonate/IPP axis. The same molecular features were previously shown to mediate selective γδ T cell killing of another pediatric cerebellar tumor, medulloblastoma, via EphA2 recognition and phosphoantigen-driven activation, while preserving normal neuronal and stem cell integrity^35^. This is clinically relevant considering separate evidence that γδ T cell abundance itself carries prognostic value in glioma: multiparametric flow cytometry across 102 GBMs and lower-grade gliomas showed that frequencies of γδ T cells and CD56^bright^ NK cells are independent positive prognostic factors^36^. Taken together, these lines of evidence suggest that the γδT-favorable sections identified here may represent rational candidates for strategies aimed at recruiting or adoptively delivering γδ T cells.

Our opportunity scoring also identified two sections (812 and 1269) as Immuno-cold and one (1239) as Suppressed, findings convergent with TIDE-based immune exclusion predictions for PFA ependymoma, which suggest that this subgroup remains immune-excluded across molecular subtypes and may require microenvironmental modulation as a prerequisite for any immunotherapy benefit^37^. The shared inhibitory program incorporated several axes with direct precedent in ependymoma or closely related pediatric CNS tumors: PD-L1/PD-L2 and IDO1 expression have been directly characterized in ependymoma immunohistochemical cohorts, COX-2 and downstream PGE synthase isoforms are among the most consistently overexpressed immunosuppressive enzymes across ependymoma and other gliomas^38,39^, and the CD39/CD73 adenosine axis has been specifically profiled in medulloblastoma, supporting its inclusion as a biologically plausible immunosuppressive mechanism in pediatric CNS tumors more broadly^40^. The HLA-E–NKG2A and Galectin-9–Tim-3 axes lack ependymoma-specific data to date but have established roles in other immune-cold pediatric (DMG) and adult (glioma) brain tumor contexts, respectively, and represent testable hypotheses for future functional validation in PFA ependymoma^41,42^. Compartment-level analysis, however, indicates that microenvironmental modulation is unlikely to be sufficient on its own, and points to which modification each modality would require. αβT ligand availability and immunosuppressive tone rose together in the mesenchymal compartment in 13 of 14 sections, so the compartment offering the most antigen-presentation machinery is also the most suppressive: any αβ-directed strategy delivers effectors into the region where suppression is highest and would need a suppression-directed partner. For γδ T cells the constraint is different. The composite γδT score showed no compartment preference, but mevalonate/IPP synthesis was higher in the epithelial compartment in 12 of 14 sections while EPHA2 and DNAM-1 ligands were higher in the mesenchymal compartment in 12 and 10 of 14. Phosphoantigen supply is therefore greatest in precisely the compartment from which lymphocyte-associated signal is most consistently absent, whereas adhesion- and ephrin-dependent recognition is best supplied where that signal is found. Transcriptional profiling reinforces this dissociation. Butyrophilin expression covaried with type I and II interferon programs in 14 of 14 sections, whereas mevalonate-pathway expression showed no reproducible interferon association and tracked lipid and glycosylation metabolism instead^43^. The two prerequisites for Vγ9Vδ2 recognition, the presenting molecules and the metabolic pathway generating their ligand, are thus neither co-regulated nor co-localized in this tumor. This argues against conditioning strategies that address only one of them: interferon-based approaches would raise butyrophilin without raising phosphoantigen supply, while aminobisphosphonate approaches would do the reverse. This provides a data-derived rationale for conditioning regimens that raise butyrophilin expression and phosphoantigen supply together, rather than either alone, as a prerequisite for γδ T cell-based approaches in this tumor. We note that these are transcript-level measures of ligand availability; BTN3A1 requires intracellular phosphoantigen binding and conformational change to signal, so transcript abundance is necessary but not sufficient.

Beyond static section-level classification, our data also point to a dynamic dimension of immunotherapy opportunity. All three matched primary–relapse pairs in our cohort (459, 723, 928) were assigned to different PFA methylation subtypes between the primary and relapse samples, consistent with the subtype instability previously documented at PFA relapse, where 13 of 32 primary-relapse pairings differed in best-matching subtype despite stable PFA-1/PFA-2 subgroup allocation^44^. Strikingly, in our data this subtype switch coincided with lymphocyte signal: all three primary sections passed lymphocyte quality control and yielded hotspot calls, whereas all three matched relapse sections fell below the detection threshold and were excluded at the screening stage. The relapse sample from patient 928 additionally showed acquisition of chromosome 1q gain, a recurrent relapse-associated copy number alteration in PFA ependymoma^14^. An important consideration is treatment exposure. All three primary sections were obtained at first resection and are treatment naive. Of the matched relapse sections, 928_2 and 723_2 were obtained after conformal radiotherapy, whereas 459_2 was not because the patient was under one year of age at diagnosis, and radiotherapy is routinely deferred in infants to avoid irradiating the developing brain. Radiotherapy is independently capable of producing both halves of the pattern we observe tumor-infiltrating lymphocytes are highly radiosensitive and are depleted locally by irradiation, while ionizing radiation simultaneously upregulates antigen-processing and presentation machinery in surviving tumor cells, including MHC class I, β2-microglobulin, TAP1/TAP2 and immunoproteasome subunits, genes that constitute the core of our αβT ligand availability program^45^. Two recent HLA ligandome studies provide the antigenic substrate for such strategies: in ependymoma specifically, analysis of naturally presented HLA class I and II ligands has identified a landscape of ependymoma-associated antigens suitable for peptide-based targeting, while a parallel atlas in medulloblastoma, a related pediatric CNS tumor, identified 15 HLA class I and 10 HLA class II frequently presented tumor-associated proteins as candidates for mRNA vaccination, peptide vaccination, dendritic cell-based approaches, or adoptive T cell transfer^46,47^. Proof-of-concept for peptide vaccination in the pediatric CNS setting already exists: a pilot trial of glioma-associated antigen (GAA) peptide vaccination in HLA-A2+ children with recurrent low-grade glioma demonstrated general tolerability and preliminary evidence of both immunological and clinical activity^48^. Section 459_2 therefore provides an informative comparison: it shows the same pattern as the irradiated relapses, with increased αβT ligand availability and lymphocyte signal below the detection threshold, despite no radiation exposure. Radiotherapy is thus not required to produce this pattern, although with a single unirradiated pair this observation is suggestive rather than conclusive, and radiation may still contribute to the two patients who received it. To our knowledge, no study has specifically examined how radiotherapy remodels the tumor immune microenvironment in ependymoma. Immune-related transcriptomic change at posterior fossa ependymoma recurrence has been documented in matched primary–relapse pairs, with sub-group-specific patterns, an immunosuppressive phenotype in PFA and increased T cell infiltration at recurrence in PFB, but treatment exposure was not disentangled from tumor evolution in that analysis^44^. Direct investigation of radiotherapy’s effect on the ependymoma TIME, ideally using matched pre- and post-irradiation spatial or single-cell transcriptomic sampling represents an important direction for future work.

Several limitations qualify these findings. Our analysis is a re-analysis of a single published Visium dataset comprising 14 sections from a small number of patients, most of them infants or toddlers, and the resulting ligand–receptor, functional state and opportunity associations will require validation in independent and larger cohorts. Cell-type and zone assignments rely on deconvolution and module scoring rather than ground-truth single-cell resolution in the same tissue. The candidate ligand–receptor axes and inhibitory program components would in particular benefit from orthogonal protein-level validation by multiplex immunofluorescence or imaging mass cytometry, as ligand–receptor scores reflect transcriptomic co-expression in spatially adjacent spots and should be read as interaction potential rather than confirmed signaling; the NicheNet analysis provides partial support for the top-ranked axes but does not substitute for experimental confirmation. A second limitation concerns resolution; lymphoid content in these sections lies close to the detection floor of 55 µm spatial transcriptomics. Canonical T cell transcripts are recovered at only a few counts per pooled region, and the inferred abundance of individual lymphoid populations did not differ statistically between hotspot and non-hotspot regions. Hotspot calls should therefore be read as regions of locally elevated aggregate lymphocyte-associated signal, robust to the choice of deconvolution reference, rather than as quantified lymphocyte infiltrates, and resolving subset composition within them will require a targeted modality such as RNAscope, multiplex immunofluorescence or region-selected profiling. For the same reason, all opportunity scores are tumor-side measurements of ligand availability and suppressive-program expression, not measurements of T cell presence. Finally, treatment exposure differs between the matched primary–relapse pairs. Two relapses followed conformal radiotherapy while the third did not, and the pairs also differ in age at first surgery and in relapse interval. The unirradiated pair argues against radiotherapy as the sole driver of the relapse-associated changes we describe, but with three pairs these contributions cannot be formally separated.

Collectively, this work provides a spatial framework for understanding immune restriction in pediatric ependymoma, and a computational strategy, reference-guided deconvolution, spatial hotspot detection, ligand–receptor inference and multi-modal T cell opportunity scoring, that can be extended to other sparsely infiltrated pediatric CNS tumors. The central biological finding is that recognition potential and lymphocyte access are governed by different, and partly opposing, features of the same tissue. These findings provide a rationale for pairing recognition-focused strategies with interventions that modify the niche associated with lymphocyte access and, for the γδ modality specifically, for conditioning approaches that jointly enhance both components of the phosphoantigen-recognition axis.

## Materials s Methods

### CIBERSORTx deconvolution and abundance calculation

Bulk RNA sequencing data from pediatric central nervous system tumors were obtained from the Open Pediatric Brain Tumor Atlas (OpenPBTA, https://portal.kidsfirstdrc.org), encompassing 19 histological entities^15^. Assessment of leukocyte fractions from the transcriptomes was performed by applying CIBERSORTx (https://cibersortx.stanford.edu/) with the LM7 immune cell signature matrix, as previously described^16,17^. Abundances were calculated from CIBERSORTx results and Sample Enrichment Scores (SES). SES was computed using the open-source software AutoCompare-SES (https://sites.google.com/site/fredsoftwares/products/autocompare_ses) with normalized settings. The open-source software DeepTIL (https://sites.google.com/site/fredsoftwares/products/deeptil) was then applied to automatically derive the abundance of seven leukocyte subsets per sample. Samples were stratified by histological diagnosis, WHO aggressiveness grade (Low, Intermediate, High, Very High), tumor anatomical location, and tumor type (primary, progression, recurrence) for downstream comparisons. Statistical comparisons across groups were performed using Kruskal-Wallis tests followed by pairwise Wilcoxon rank-sum tests with Benjamini–Hochberg correction, implemented in the rstatix R package (v0.7.3).

### Semla spatial deconvolution

Spatial deconvolution was performed using the non-negative least squares (NNLS) implementation provided by Semla (v1.4.0; RunNNLS, https://github.com/spatial-research/semla). The scRNA-seq reference was derived from the pediatric ependymoma cohort GSE125969 and consisted of an immune-focused subset containing the major immune cell populations together with limited representation of neoplastic cell states. All cell-type groups present in this reduced reference were included simultaneously in the NNLS deconvolution; the reference was not subset separately for individual spatial sections. This design was chosen to retain competing non-lymphocyte programs while maintaining sensitivity to the relatively sparse immune signal in PFA ependymoma.

The reference object was constructed in Seurat (v5.3.1) using cells passing minimum quality thresholds (≥3 cells per gene, ≥200 genes per cell), followed by library-size normalization, identification of highly variable features, data scaling, and PCA. Each spatial section was processed independently using relative count normalization (scale factor 10,000), selection of the top 3,000 highly variable features, scaling, and PCA prior to deconvolution. Only the Lymphocyte NNLS coefficient was carried forward for hotspot detection and downstream spatial analyses. To assess whether limited representation of neoplastic states in the immune-focused reference could artificially inflate the lymphocyte signal, the NNLS analysis was repeated using a tumor-augmented reference in which additional PFA tumor cells were added to the original reference. Concordance between the original and tumor-augmented analyses was evaluated as a sensitivity analysis of hotspot identification.

For each of the 14 sections, three quality-control metrics were calculated from the per-spot Lymphocyte NNLS coefficients: the number of spots with non-missing values (≥50), the proportion of spots with a non-zero coefficient (≥0.10), and the maximum observed coefficient (≥0.05). Seven sections failed the automated screen, all on the non-zero proportion criterion. Because these criteria are lower bounds, they cannot identify sections with globally inflated lymphocyte coefficients. The seven passing sections were therefore additionally inspected spatially. Section 848 was excluded because lymphocyte signal was distributed near-uniformly across the capture area rather than forming spatially restricted foci. The remaining six sections (459, 723, 812, 821, 928, and 1239) were retained for downstream hotspot analysis.

### Lymphocyte hotspot detection

Lymphocyte infiltration hotspots were identified within tissue-only spots (as defined by the in_tissue flag from the Visium tissue position file) using a two-criterion spatial scoring approach. For each tissue spot, a local neighborhood score was computed as the mean Lymphocyte NNLS score across the spot itself and its six nearest neighbors in high-resolution pixel coordinate space (k-nearest neighbor graph, k = 6), computed using the dbscan R package (v1.2.3). A spot was classified as a hotspot if it simultaneously exceeded the 90th percentile of the section-level Lymphocyte NNLS-coefficient distribution and the 90th percentile of the local neighborhood score distribution. This dual threshold was designed to identify spatially coherent high-infiltration foci rather than isolated high-scoring spots.

### Spatial pseudobulk generation and immune abundance calculation

To characterize the immune composition of Semla-defined lymphocyte hotspots, section-specific spatial pseudobulk expression profiles were generated from the Visium data. For each tissue section, raw gene counts were summed independently across spots classified as lymphocyte hotspots and across the remaining in-tissue non-hotspot spots, generating paired hotspot and non-hotspot pseudobulk profiles. Pseudobulks were generated separately for each tissue section to preserve section-level biological replication and were not pooled across samples.

Immune cell fractions were estimated from the hotspot and non-hotspot pseudobulks by applying CIBERSORTx with the LM7 immune cell signature matrix, using the same workflow as for the bulk RNA-sequencing analysis described above. Because LM7-derived CIBERSORTx fractions represent the relative composition of the modeled leukocyte compartment, immune cell abundances were additionally calculated using Sample Enrichment Scores (SES) and DeepTIL. SES was computed using AutoCompare-SES with normalized settings, and DeepTIL was applied to derive abundance estimates for the seven LM7 leukocyte subsets. Hotspot and non-hotspot estimates were compared within each tissue section.

### Tumor region zone annotation

Spatial zone programs were defined for each of the six lymphocyte hotspot-positive sections using marker gene sets derived from initial Visium data cluster annotations of PFA ependymoma^14^. Four biologically meaningful zones were defined by aggregating cluster-level marker genes into the following groups: Epithelial (clusters TEC-A, TEC-B, TEC-C, TEC-D, UEC-A, CEC), Mesenchymal (clusters MEC-A, MEC-B, MEC-C, MEC-D, UEC-B), Vascular (cluster VE), and Myeloid (clusters classic-M, hypoxia-M, chemokine-M). For each cluster, marker genes passing the Δpct ≥ 0.15 filter were ranked first by adjusted *P* value and subsequently by a composite specificity score defined as avg_log2FC × Δpct, with the top 100 genes per cluster retained. For the Vascular zone, additional filters were applied (pct.1 ≥ 0.20, avg_log2FC ≥ 0.40, pct.2 ≤ 0.30), and the resulting gene set was further restricted to the intersection with a curated list of 22 canonical endothelial marker genes (including *PECAM1, VWF, KDR, FLT1, CD34, EMCN, PLVAP*, and related markers) to avoid contamination by non-endothelial signals. After union across constituent clusters, gene symbols were converted to uppercase and mapped to the Visium gene space of each section.

Zone-specific module scores were computed per Visium spot using AddModuleScore from the Seurat package (v 5.3.1). Scores were z-scored within each section to correct inter-section baseline differences. Each in-tissue spot was assigned to a dominant zone label corresponding to the zone with the highest z-scored module score, subject to a minimum z-score threshold of 0.5. Spots for which no zone program exceeded this threshold were labeled “Uncertain.”

### Permutation-based spatial neighborhood analysis

To validate zone associations at single-spot resolution and account for section-level differences in global zone composition, a spatial neighborhood permutation test was performed independently for each hotspot-positive section. For each in-tissue spot with valid pixel coordinates, the six nearest neighbors (6-NN) were identified using Euclidean distance in high-resolution pixel coordinate space (pxl_col_in_hires, pxl_row_in_hires), computed using the kNN function from the dbscan R package (v 1.2.3). For each spot, the fraction of its 6 nearest neighbors belonging to each of the Epithelial, Myeloid, Mesenchymal, and Vascular zones was recorded as a neighbor’s fraction score.

The observed test statistic was defined as the mean neighbor fraction score across all hotspot-positive spots within a section. A null distribution was generated by 1,000 random permutations: in each permutation, hotspot labels were randomly reassigned to the same number of in-tissue spots (without replacement), and the mean neighbor fraction was recomputed. Empirical p-values were calculated as:

For enrichment tests (Myeloid, Mesenchymal): p = (number of permutations ≥ observed + 1) / (1,000 + 1)

For depletion tests (Epithelial): p = (number of permutations ≤ observed + 1) / (1,000 + 1)

P values were adjusted within each test (epithelial depletion, myeloid enrichment, mesenchymal enrichment, vascular enrichment, interface enrichment) across the six hotspot-positive sections using the Benjamini–Hochberg procedure. All analyses used a fixed random seed (set.seed(1)). Sections with fewer than 5 hotspot spots were excluded from permutation testing. Statistical significance was reported using the same thresholds as above.

### Signature independence analysis

To confirm that the spatial co-localization of lymphocyte hotspots with mesenchymal and myeloid zones does not reflect shared gene measurement artefacts, we assessed pairwise overlap between the three zone-defining gene signatures. The lymphocyte gene set was derived independently from the same snRNA-seq reference using FindMarkers (Seurat), comparing the Lymphocyte cluster against all other cell types, and retaining the top 200 most discriminative marker genes by adjusted p-value and log2 fold change. The Mesenchymal (306 genes) and Myeloid (182 genes) gene sets were identical to those used for zone scoring. Pairwise overlaps were quantified by intersection size, percentage of each gene set represented in the overlap, and Jaccard index (|A ∩ B| / |A ∪ B|).

### Lymphocyte hotspot functional state scoring

Functional state programs were scored in all in-tissue spots using AddModuleScore_UCell from the UCell R package, applied independently to each Visium section^49^. Five curated gene sets were used, each designed to capture a distinct T cell or innate-like lymphocyte functional state:

<u>Cytotoxic</u>: *NKG7, PRF1, GZMB, GNLY, CTSW, KLRD1, KLRK1, XCL1, XCL2, CCL5, IFNG*

<u>Inflammatory</u>: *IFNG, TNF, CCL4, CCL5, XCL1, XCL2, NFKBIA, IRF1, STAT1*

<u>Exhausted:</u> *PDCD1, LAG3, TIGIT, HAVCR2, CTLA4, TOX, ENTPD1, CXCL13*

<u>IL17-like:</u> *RORC, CCR6, KLRB1, IL7R, IL17A, IL17F, AREG*

<u>Retention:</u> *CD44, ITGAE, CXCR3, CXCRc, CDcS, CCL5*

For each hotspot, functional-state assignment was restricted to signatures with sufficient transcriptional support to avoid forced classification of weakly matching programs. UCell scores were calibrated separately for each signature and section relative to the distribution of that signature across lymphocyte hotspot spots. A signature was considered eligible when its score was at or above the 50th percentile of the corresponding section-specific hotspot distribution and at least two genes from the signature were detected in the individual spot. When multiple signatures fulfilled these criteria, the program with the highest percentile-normalized score was assigned as the dominant functional state only when it exceeded the second-ranked eligible program by at least 0.05 on the percentile-normalized 0–1 scale (five percentile points). Hotspots not satisfying these criteria were classified as unassigned.

Gene sets were matched to genes present in the corresponding Visium expression matrix before scoring, and signatures were evaluated using the remaining detected genes when individual signature genes were absent. Unassigned hotspots were retained in analysis outputs and quality-control summaries but excluded when calculating the relative functional-state composition of confidently classified hotspots shown in Fig. 4b.

### Spatial ligand-receptor interaction scoring

Candidate ligand–receptor (LR) pairs were independently drawn from two curated resources: CellChatDB (human), obtained from the CellChat repository using a fixed repository commit, and CellPhoneDB, accessed via liana::select_resource(“CellPhoneDB”) from the LIANA R package (v0.1.14)^50,51^. Both resources were filtered to retain only interactions for which both ligand and receptor genes were detectable across the union of all section expression matrices, using a presence-based pre-filter applied per entity, including individual genes and multi-subunit receptor complexes.

For each resource independently, LR interaction scores were calculated per section and spatial zone using the same spatial scoring framework. A directed 6-nearest-neighbour (6-NN) spatial graph was constructed within each section using Euclidean distance in high-resolution pixel-coordinate space. Spots assigned to the Myeloid, Mesenchymal or Vascular zones were treated as sender spots, whereas lymphocyte-hotspot spots were treated as receivers. For each LR pair and section, an edge-level interaction score was calculated for each zone-to-hotspot spatial edge as the product of ligand expression in the sender, receptor expression in the receiver, the non-negative sender zone-module score, and the receiver’s rescaled Lymphocyte NNLS coefficient. The section-level interaction score was defined as the mean across eligible edges. Across sections, each interaction was summarized by the number of supporting sections and its mean, median, and maximum score. Myeloid and Mesenchymal interactions were ranked first by the number of supporting sections, followed by median and maximum score, whereas Vascular interactions were ranked by maximum score, followed by section support and median score. The top 30 interactions per zone were retained independently for each LR resource. Five sections displaying the recurrent hotspot-associated niche pattern (459, 812, 821, 928 and 1239) were included in this analysis. Section 723 was excluded because it did not display the recurrent neighborhood pattern observed in the other hotspot-positive sections. Section 1239 was additionally excluded from the Myeloid analysis and section 928 from the Vascular analysis because of insufficient zone-specific spatial support. Statistical robustness was assessed using 1,000 hotspot-label permutations per zone and section, with randomization controlled using a fixed seed. Empirical permutation *P* values were used as a robustness measure and were not used to define the top-30 interaction lists.

A joint shortlist of LR pairs was constructed by joining the CellChatDB and CellPhoneDB top-30 ranked lists within each zone on ligand–receptor identity. Interactions were annotated according to whether they occurred among the CellChatDB top-ranked pairs, the CellPhoneDB top-ranked pairs, or both. For cross-database shared pairs, min_score_both was defined as the minimum of the two resource-specific mean interaction scores. In Fig. 4f, fill represents min_score_both, whereas point size represents cross-database section support as a percentage of eligible sections.

### Cross-section quality control of ligand–receptor interactions

Cross-section quality control was performed independently for each LR resource and spatial zone to determine whether aggregate interaction patterns were disproportionately driven by individual sections. At the section level, the total number of detected LR pairs and the mean, median and maximum observed interaction scores were calculated. The mean interaction score and number of detected pairs were standardized across sections within each resource–zone combination, and sections with an absolute standardized value ≥2 were flagged as potential outliers.

For top-ranked LR pairs, section-specific interaction scores were additionally examined to assess cross-section consistency. For each interaction, the number of supporting sections, mean, standard deviation, coefficient of variation, and minimum and maximum interaction scores were calculated. Pair-specific scores were standardized across sections, and section–interaction combinations with an absolute standardized score ≥2 were flagged for inspection.

Permutation results were also evaluated at the section level. For each resource, zone and section, the number and fraction of tested interactions with empirical *P* < 0.05 and *P* < 0.01 were calculated. The fraction of interactions reaching *P* < 0.05 was standardized across sections within each resource–zone combination, and sections with an absolute standardized value ≥2 were flagged as potential outliers.

### NicheNet ligand activity and downstream target gene prediction

NicheNet analysis was used to evaluate whether candidate ligands identified by the spatial LR analysis could account for transcriptional changes in spatially adjacent lymphocyte hotspots. Human NicheNet prior networks were used, including the ligand–target regulatory potential matrix (ligand_target_matrix_nsga2r_final.rds), ligand–receptor network (lr_network_human_21122021.rds) and weighted signaling network (weighted_networks_nsga2r_final.rds)^52^. For each evaluated spatial interface, hotspot spots with at least one of their six nearest neighbors assigned to the corresponding zone were classified as zone-adjacent (“touch”), whereas hotspot spots lacking a neighbor from that zone were classified as “not_touch”. Differential expression between touch and not_touch hotspot groups was calculated independently within each eligible section using FindMarkers in Seurat with the Wilcoxon rank-sum test and a minimum detection fraction of 0.05. Sections were excluded when either group contained fewer than 15 hotspot spots. Genes with adjusted *P* < 0.20 and average log2 fold-change >0.10 were retained as genes upregulated in zone-adjacent hotspots.

Section-specific upregulated genes were aggregated across eligible sections to generate a cross-section receiver gene set. For each gene, the number of sections in which it was upregulated and its mean log2 fold-change were calculated. Genes were ranked first by the number of supporting sections and subsequently by mean log2 fold-change; genes supported by at least one section were eligible for inclusion, and up to the top 300 ranked genes were retained. Thus, recurrence across sections was used as a ranking criterion rather than as a formal consensus requirement. Candidate ligands were obtained from the union of ligands represented among the top CellChatDB and CellPhoneDB interactions for the corresponding spatial zone. Candidate ligands were restricted to those represented in the NicheNet ligand–target matrix. Background genes were defined as genes represented in the ligand–target matrix and detected in the spatial transcriptomic dataset.

For each candidate ligand, NicheNet ligand–target regulatory potential scores were evaluated across the background gene set. Ligand activity was quantified using an area-under-the-curve (AUC) statistic measuring whether genes in the receiver gene set preferentially received higher predicted regulatory potential than the remaining background genes. Candidate ligands were ranked according to their AUC. In addition to the cross-section analysis, NicheNet ligand activity was calculated independently within each eligible section using that section’s own set of genes upregulated in zone-adjacent hotspots. Section-level ligand activity was calculated only when at least 15 receiver genes were available and was retained to evaluate the reproducibility of predicted ligand activity across spatial sections.

### Immunotherapy opportunity scoring

Immunotherapy opportunity scores were computed per Visium spot across all 14 ependymoma sections, restricted to tumor-cell zones (Epithelial and Mesenchymal) as assigned above. Three gene programs were defined a priori and applied unchanged to every section:

αβT core opportunity (12 genes): HLA-A, HLA-B, HLA-C, B2M, TAP1, TAP2, PSMB8, PSMB9, ERAP1, ERAP2, NLRC5, IRF1 — MHC class I antigen presentation machinery and immunoproteasome components.

γδT ligand availability (17 genes): BTN2A1, BTN3A1; MVK, PMVK, MVD, IDI1; MICA, MICB, ULBP2, ULBP3, RAET1E, RAET1G, RAET1L; EPHA2; PVR, NECTIN2, ICAM1 — butyrophilins, mevalonate pathway, NKG2D stress ligands, ephrin receptor, and DNAM-1 ligands and adhesion molecules.

Immunosuppressive tone (16 genes): CD274, PDCD1LG2, HLA-E, LGALS9, IDO1, TGFB1, TGFB2, TGFB3, IL10, VEGFA, NT5E, ENTPD1, PTGS2, PTGES, PTGER2, PTGER4 — checkpoint ligands, immunosuppressive cytokines, adenosine-pathway ectoenzymes and COX–PGE2 components.

Genes detected in fewer than 5% of tumor spots were excluded from scoring; 12 of 12 αβT, 11 of 17 γδT and 9 of 16 immunosuppressive genes were retained. Expression was log-normalized per section and each gene z-transformed across the pooled set of tumor spots from all sections; scores are therefore relative to this 14-section cohort and are not absolute measures of ligand availability or immunosuppression.

Gene z-scores were averaged within predefined mechanism axes; axis scores were z-transformed and averaged with equal weight to give each composite score, which was z-transformed in turn. The axes were butyrophilin (BTN2A1, BTN3A1), mevalonate (MVK, PMVK, MVD, IDI1), NKG2D ligand (MICA), ephrin (EPHA2) and DNAM-1/adhesion (PVR, NECTIN2, ICAM1) for the γδT program; and HLA-E, galectin-9 (LGALS9), TGF-β (TGFB1–3), adenosine (NT5E, ENTPD1), COX-2 (PTGS2) and VEGF (VEGFA) for the immunosuppressive program. The IL-10 axis retained no gene above the detection threshold. The αβT core was not partitioned. After detection filtering, several axes retained a single gene (NKG2D ligand, MICA; ephrin, EPHA2; COX-2, PTGS2; VEGF, VEGFA); axes were weighted equally regardless of the number of genes retained, so single-gene axes contribute as much as multi-gene axes to the composite.

Net scores were computed spot-wise as αβT_net = αβT_core_z − immunosuppressive_z and γδT_net = γδT_core_z − immunosuppressive_z and summarized as the median across in-tissue tumor spots per section. Sections were assigned to five opportunity categories: Dual-opportunity (both nets > 0), αβT-favorable (αβT_net > 0 and γδT_net ≤ 0), γδT-favorable (αβT_net ≤ 0 and γδT_net > 0), Suppressed (both ≤ 0 and median immunosuppressive z ≥ 0.75), and Immuno-cold (all remaining). Categories are a discretization of continuous scores, and sections close to a boundary should be interpreted accordingly. Mechanism axes were summarized per section using the mean rather than the median, because for sparsely detected axes the section median reduces to the value of zero on the z scale.

### Scoring quality control

Split-half analysis (spots randomly halved within each section, 100 repetitions) gave section-level Spearman ρ = 0.956 (αβT net) and 0.952 (γδT net); category assignment agreed between halves in 81.6% of section-repetition pairs. Classification agreed in 78.6–92.9% of sections across gene-wise z-mean, raw-mean and principal-component aggregation. Rank-based scoring (UCell) was evaluated and not adopted: it coupled the αβT and γδT net scores to sequencing depth to different degrees (ρ = 0.24 and −0.02) whereas z-mean scoring coupled both approximately equally (ρ = 0.15 and 0.13). Sequencing depth did not account for section-level differences (ρ = 0.18 and −0.08, n = 14). Classification was unchanged in all 14 sections when the mevalonate genes were removed from the γδT program.

### Compartment and spatial analysis

Opportunity scores were compared between the epithelial and mesenchymal tumor compartments defined above. Per-spot scores were summarized as the section median within each compartment (mechanism axes: section mean) and compared using a two-sided paired Wilcoxon signed-rank test across the 14 sections, with the section as the unit of replication; spot-level testing was avoided because spots within a section are not independent. P values were adjusted by the Benjamini–Hochberg method separately within each family of tests: the three opportunity components, the six γδT recognition modalities, the six immunosuppressive mechanisms, and the two net scores. The number of sections in which each score was higher in the epithelial compartment is reported alongside the adjusted p value. As a sensitivity analysis, each score was regressed on log sequencing depth within section and the residuals summarized and tested identically; between-section depth differences were retained deliberately, being confounded with patient. As a further sensitivity analysis addressing non-independence between matched primary–relapse sections, compartment comparisons for the three opportunity components were repeated using one section per patient (n = 11, relapse sections excluded). The significance pattern was unchanged in the patient-independent analysis: αβT ligand availability and immunosuppressive tone remained significantly higher in the mesenchymal compartment, whereas γδT ligand availability remained non-significantly different between compartments.

Recognition-modality balance was defined per spot as γδT net − αβT net. Because both net scores subtract the same immunosuppressive term, this contrast is independent of inhibitory tone. Balance values were mapped onto high-resolution tissue coordinates and overlaid on the corresponding HCE image, with color limits shared across sections and clipped at the 2nd and 98th percentiles.

### Gene set enrichment analysis of opportunity scores

To identify transcriptional programs associated with the T-cell opportunity landscape while minimizing section-or patient-specific effects, gene set enrichment analysis (GSEA) was performed from within-section gene–score associations. For each spatial transcriptomic section, Spearman correlations were calculated between normalized expression of each gene and each spot-level opportunity score across tumor spots. Genes were required to be detected in at least 10% of tumor spots within a section and to be evaluable in at least 10 sections. For each gene, section-specific Spearman correlation coefficients were Fisher *z*-transformed and integrated across sections using a weighted Stouffer method, with weights proportional to (\sqrt{n-3}), where *n* represents the number of informative spots in the corresponding section. The resulting combined *z*-statistics provided a cross-section ranking of genes according to their positive or negative association with each score. Ribosomal protein, mitochondrial and hemoglobin genes were excluded from the enrichment rankings to reduce compositional effects associated with library-size normalization. Their gene–score correlations were nevertheless retained and evaluated separately as an analysis diagnostic.

GSEA was performed on the combined gene rankings using fgseaMultilevel from the fgsea R package. Gene sets were obtained from MSigDB using msigdbr and included the Hallmark and Reactome collections. Gene sets containing 10–500 genes were tested, and pathways with Benjamini–Hochberg-adjusted *P* < 0.05 were considered significantly enriched. Redundant significant pathways were reduced using collapsePathways, with representative main pathways retained for interpretation and visualization. To assess the reproducibility of pathway associations across tumors, GSEA was additionally performed independently for each spatial section using the corresponding section-specific gene–score correlations. For each pathway, cross-section concordance was quantified as the number of sections in which the normalized enrichment score (NES) had the same direction as the combined analysis. Genes constituting the opportunity scores were retained in the ranked gene lists. Enriched pathways were therefore additionally annotated according to whether they contained genes contributing to the corresponding score, allowing associations overlapping the scoring program to be distinguished from pathway associations independent of the score definition.

## Author contributions (with CRediT role information)

**P.T.**: Conceptualization, Methodology, Software, Formal analysis, Visualization, Writing – review & editing. **R.M.**: Conceptualization, Writing – review & editing. **J.L**.: Supervision, Writing – review & editing. **K.B**.: Conceptualization, Writing – review & editing, Supervision, Funding acquisition. **L.B**.: Conceptualization, Methodology, Formal analysis, Investigation, Writing – original draft, Writing – review & editing, Visualization, Project administration.

## Ethics approval

This study is a secondary computational analysis of publicly available, de-identified data deposited in the Gene Expression Omnibus (GSE195661 and GSE125969). No new patient material was collected, no new patient data were generated, and no direct patient contact occurred.

## Data availability

Visium spatial transcriptomics data are available from the Gene Expression Omnibus under accession GSE195661. The scRNA-seq reference dataset used for deconvolution is available under accession GSE125969. Bulk RNA-sequencing data used for the pan-pediatric CNS tumor analysis were obtained from the Open Pediatric Brain Tumor Atlas (OpenPBTA). Code supporting the analyses presented in this study is available at https://github.com/Lola-Boutin/epn-spatial-immune.”

## References

1. Korshunov, A. et al. Molecular staging of intracranial ependymoma in children and adults. J. Clin. Oncol. Off. J. Am. Soc. Clin. Oncol. 28, 3182–3190 (2010).

2. Duffner, P. K. et al. Prognostic Factors in Infants and Very Young Children with Intracranial Ependymomas. Pediatr. Neurosurg. 28, 215–222 (1998).

3. Jagtiani, P. et al. Intracranial ependymomas in pediatric patients: patterns of care, disparities, and survival outcomes from the National Cancer Database. J. Neurosurg. Pediatr. 34, 495–508 (2024).

4. Yu, Q.-S., Yin, Y.-H. & Yu, X.-G. Clinical Characteristics, Treatment, and Survival Outcome of Ependymoma in Infants. World Neurosurg. 181, e75–e83 (2024).

5. Pajtler, K. W. et al. Molecular Classification of Ependymal Tumors across All CNS Compartments, Histopathological Grades, and Age Groups. Cancer Cell 27, 728–743 (2015).

6. Kresbach, C., Neyazi, S. & Schüller, U. Updates in the classification of ependymal neoplasms: The 2021 WHO Classification and beyond. Brain Pathol. 32, e13068 (2022).

7. Pohl, L. C. et al. Molecular characteristics and improved survival prediction in a cohort of 2023 ependymomas. Acta Neuropathol. (Berl.) 147, 24 (2024).

8. Gröbner, S. N. et al. The landscape of genomic alterations across childhood cancers. Nature 555, 321–327 (2018).

9. Nabbi, A. et al. Transcriptional immunogenomic analysis reveals distinct immunological clusters in paediatric nervous system tumours. Genome Med. 15, 67 (2023).

10. Galvin, R. T., Jena, S., Maeser, D., Gruener, R. & Huang, R. S. Revealing Pan-Histology Immunomodulatory Targets in Pediatric Central Nervous System Tumors. Cancers 15, 5455 (2023).

11. Hwang, E. I. et al. The current landscape of immunotherapy for pediatric brain tumors. *Nat*. Cancer 3, 11–24 (2022).

12. Grabovska, Y. et al. Pediatric pan-central nervous system tumor analysis of immune-cell infiltration identifies correlates of antitumor immunity. Nat. Commun. 11, 4324 (2020).

13. Wu, H. et al. Single-Cell RNA Sequencing Unravels Upregulation of Immune Cell Crosstalk in Relapsed Pediatric Ependymoma. Front. Immunol. 13, 903246 (2022).

14. Fu, R. et al. Spatial transcriptomic analysis delineates epithelial and mesenchymal subpopulations and transition stages in childhood ependymoma. Neuro-Oncol. 25, 786–798 (2023).

15. Shapiro, J. A. et al. OpenPBTA: The Open Pediatric Brain Tumor Atlas. Cell Genomics 3, 100340 (2023).

16. Newman, A. M. et al. Determining cell type abundance and expression from bulk tissues with digital cytometry. Nat. Biotechnol. 37, 773–782 (2019).

17. Tosolini, M. et al. Assessment of tumor-infiltrating TCRVγ9Vδ2 γδ lymphocyte abundance by deconvolution of human cancers microarrays. Oncoimmunology 6, e1284723 (2017).

18. Turner, C. P. et al. Tumour infiltrating lymphocyte density differs by meningioma type and is associated with prognosis in atypical meningioma. Pathology (Phila.) 54, 417–424 (2022).

19. Rapp, C. et al. Cytotoxic T Cells and their Activation Status are Independent Prognostic Markers in Meningiomas. Clin. Cancer Res. 25, 5260–5270 (2019).

20. Lieberman, N. A. P. et al. Characterization of the immune microenvironment of diffuse intrinsic pontine glioma: implications for development of immunotherapy. Neuro-Oncol. 21, 83–94 (2019).

21. Zhang, Q. et al. Interrogation of the microenvironmental landscape in spinal ependymomas reveals dual functions of tumor-associated macrophages. Nat. Commun. 12, 6867 (2021).

22. Griesinger, A. M. et al. Multi-omic approach identifies hypoxic tumor-associated myeloid cells that drive immunobiology of high-risk pediatric ependymoma. iScience 26, 107585 (2023).

23. Larsson, L., Franzén, L., Ståhl, P. L. & Lundeberg, J. Semla: a versatile toolkit for spatially resolved transcriptomics analysis and visualization. Bioinformatics 39, btad626 (2023).

24. Savage, N. et al. Dual tumour–myeloid targeting of glioblastoma with GPNMB CAR-T cells. Nature 1–10 (2026) doi:10.1038/s41586-026-10641-1.

25. Jhunjhunwala, S., Hammer, C. & Delamarre, L. Antigen presentation in cancer: insights into tumour immunogenicity and immune evasion. Nat. Rev. Cancer 21, 298–312 (2021).

26. Sebestyen, Z., Prinz, I., Déchanet-Merville, J., Silva-Santos, B. & Kuball, J. Translating gammadelta (γδ) T cells and their receptors into cancer cell therapies. Nat. Rev. Drug Discov. **1G**, 169–184 (2020).

27. Zhang, J. et al. Single-cell and spatial transcriptomics reveal SPP1-CD44 signaling drives primary resistance to immune checkpoint inhibitors in RCC. J. Transl. Med. 22, 1157 (2024).

28. Khan, K. A. et al. Modulation of fibronectin extracellular matrix enhances anti-tumor efficacy of immune checkpoint blockade. Cell Rep. Med. 6, 102322 (2025).

29. Vijver, S. V. et al. Collagen Fragments Produced in Cancer Mediate T Cell Suppression Through Leukocyte-Associated Immunoglobulin-Like Receptor 1. Front. Immunol. 12, (2021).

30. Ma, C. et al. Therapeutic modulation of APP-CD74 axis can activate phagocytosis of TAMs in GBM. Biochim. Biophys. Acta Mol. Basis Dis. 1870, 167449 (2024).

31. Porter, T. R., Inyushin, M. & Kucheryavykh, L. Amyloid precursor protein accumulation in glioblastoma is associated with altered synaptic dynamics and immune suppression. *Discov*. Oncol. 16, 1730 (2025).

32. Gielen, P. R. et al. Elevated levels of polymorphonuclear myeloid-derived suppressor cells in patients with glioblastoma highly express S100A8/9 and arginase and suppress T cell function. Neuro-Oncol. 18, 1253–1264 (2016).

33. Cómitre-Mariano, B. et al. S100A proteins show a spatial distribution of inflammation associated with the glioblastoma microenvironment architecture. Theranostics 15, 726–744 (2025).

34. Zhang, D., Ma, Y. & Feng, D. Mapping TAM–tumor crosstalk in glioma via ligand–receptor multi-omics: mechanisms of immune evasion. Front. Immunol. 16, (2025).

35. Boutin, L., Liu, M., Déchanet Merville, J., Bedoya-Reina, O & Wilhelm, M. T. EphA2 and phosphoantigen-mediated selective killing of medulloblastoma by γδT cells preserves neuronal and stem cell integrity. OncoImmunology 14, 2485535 (2025).

36. Gershon, R. et al. Frequencies of 4 tumor-infiltrating lymphocytes potently predict survival in glioblastoma, an immune desert. Neuro-Oncol. 26, 473–487 (2024).

37. Palermo, M. et al. Algorithm-based assessment of T-cell dysfunction and exclusion to forecast ICB sensitivity in pediatric brain ependymoma. J. Neurooncol. 176, 128 (2025).

38. Kim, S.-K. et al. Overexpression of cyclooxygenase-2 in childhood ependymomas: role of COX-2 inhibitor in growth and multi-drug resistance in vitro. Oncol. Rep. 12, 403–409 (2004).

39. Baryawno, N. et al. Tumor-growth-promoting cyclooxygenase-2 prostaglandin E2 pathway provides medulloblastoma therapeutic targets. Neuro-Oncol. 10, 661–674 (2008).

40. Stefani, M. A. et al. ENTPD1 (CD39) and NT5E (CD73) expression in human medulloblastoma: an in silico analysis. Purinergic Signal. 21, 331–337 (2025).

41. Deng, Y. et al. Targeting the HLA-E-NKG2A axis in combination with MS-275 enhances NK cell-based immunotherapy against DMG. J. Exp. Clin. Cancer Res. CR 44, 133 (2025).

42. Lee, C. et al. Galectin-9 Mediates the Functions of Microglia in the Hypoxic Brain Tumor Microenvironment. Cancer Res. 84, 3788–3802 (2024).

43. Seo, M. et al. MAP4-regulated dynein-dependent trafficking of BTN3A1 controls the TBK1–IRF3 signaling axis. Proc. Natl. Acad. Sci. U. S. A. 113, 14390–14395 (2016).

44. Hoffman, L. M. et al. Molecular sub-group-specific immunophenotypic changes are associated with outcome in recurrent posterior fossa ependymoma. Acta Neuropathol. (Berl.) 127, 731–745 (2014).

45. Zhu, S., Wang, Y., Tang, J. & Cao, M. Radiotherapy induced immunogenic cell death by remodeling tumor immune microenvironment. Front. Immunol. 13, 1074477 (2022).

46. Mühlenbruch, L. et al. The immunopeptidomic landscape of ependymomas provides actionable antigens for T-cell-based immunotherapy. Neuro-Oncol. Adv. 7, vdae226 (2025).

47. Velz, J. et al. Mapping naturally presented T cell antigens in medulloblastoma based on integrative multi-omics. Nat. Commun. 16, 1364 (2025).

48. Pollack, I. F. et al. Immune responses and outcome after vaccination with glioma-associated antigen peptides and poly-ICLC in a pilot study for pediatric recurrent low-grade gliomas. Neuro-Oncol. 18, 1157–1168 (2016).

49. Andreatta, M. & Carmona, S. J. UCell: Robust and scalable single-cell gene signature scoring. Comput. Struct. Biotechnol. J. 19, 3796–3798 (2021).

50. Efremova, M., Vento-Tormo, M., Teichmann, S. A. & Vento-Tormo, R. CellPhoneDB: inferring cell–cell communication from combined expression of multi-subunit ligand–receptor complexes. Nat. Protoc. 15, 1484–1506 (2020).

51. Jin, S. et al. Inference and analysis of cell-cell communication using CellChat. Nat. Commun. 12, 1088 (2021).

52. Browaeys, R., Saelens, W. & Saeys, Y. NicheNet: modeling intercellular communication by linking ligands to target genes. Nat. Methods 17, 159–162 (2020).

