## Supplementary figures for "Spatial immune profiling reveals lymphocyte confinement to myeloid–mesenchymal niches and heterogeneous therapeutic T-cell opportunity in pediatric ependymoma"

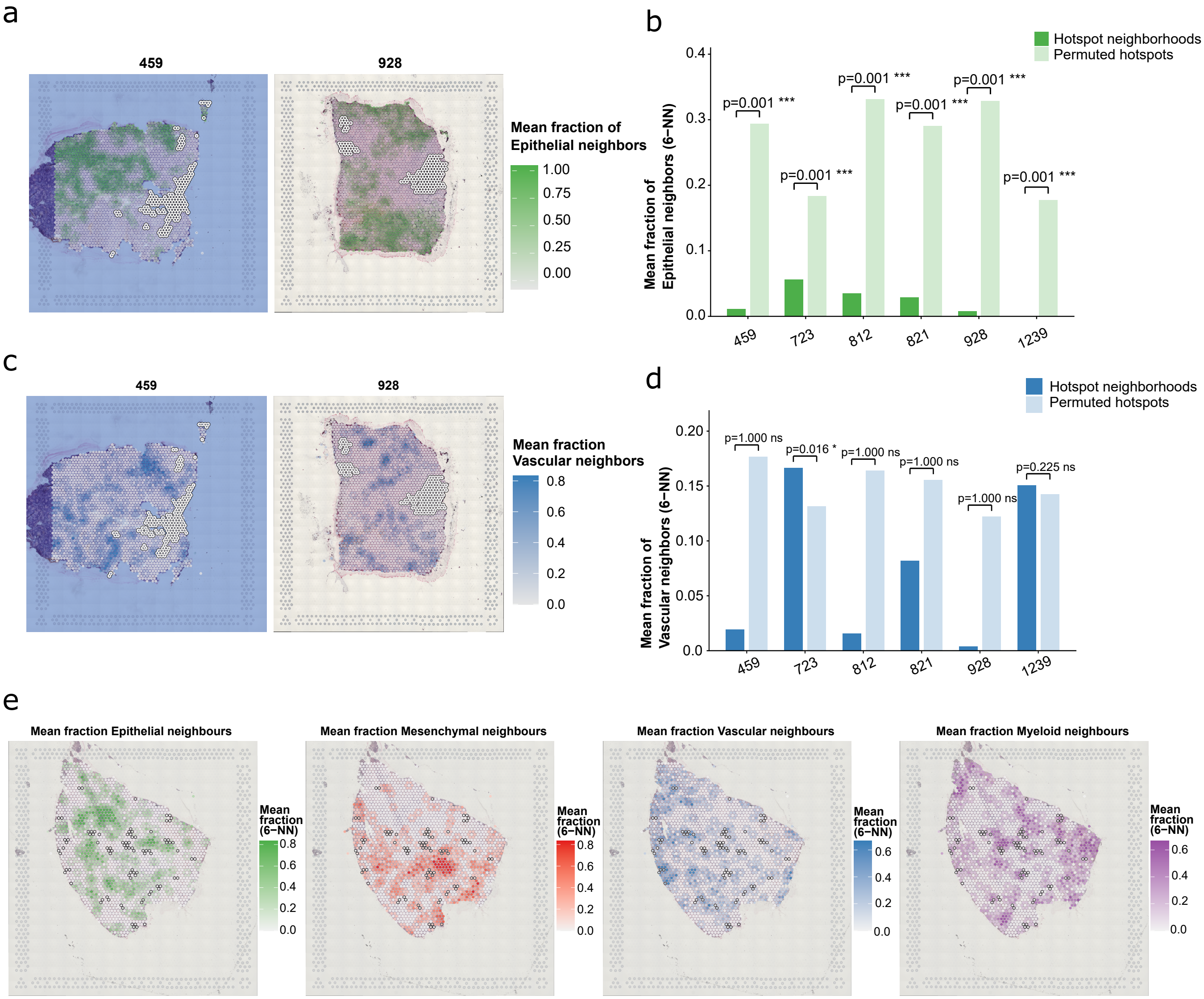


Fig. S1: Neighborhood analysis of Epithelial and Vascular zones, and atypical zone confinement pattern in section 723.
(a) Spatial maps of mean fraction of Epithelial neighbors (6-nearest-neighbour radius) per spot for sections 459 and 928, illustrating depletion of epithelial context in and around lymphocyte hotspot regions.
(b) Bar chart comparing the mean fraction of Epithelial neighbors in observed hotspot spots (dark green) versus 1,000 permuted hotspot labeling (light green) across six hotspot-positive sections. Hotspot spots are consistently and significantly depleted of Epithelial neighbors relative to permuted expectation across all six sections. Significance was assessed using empirical permutation P values followed by Benjamini–Hochberg correction; *** BH-adjusted $P < 0.001$.
(c) Spatial maps of mean fraction of Vascular neighbors per spot for sections 459 and 928.
(d) Bar chart comparing observed versus permuted mean fraction of Vascular neighbors across six hotspot-positive sections. No consistent enrichment of vascular context is observed in hotspot spots, arguing against a perivascular infiltration model. Significance was assessed using empirical permutation P values followed by Benjamini–Hochberg correction; * BH-adjusted $P < 0.05$.
(e) Zone annotation map for section 723, the only hotspot-positive section lacking consistent zone confinement in neighborhood permutation analyses.

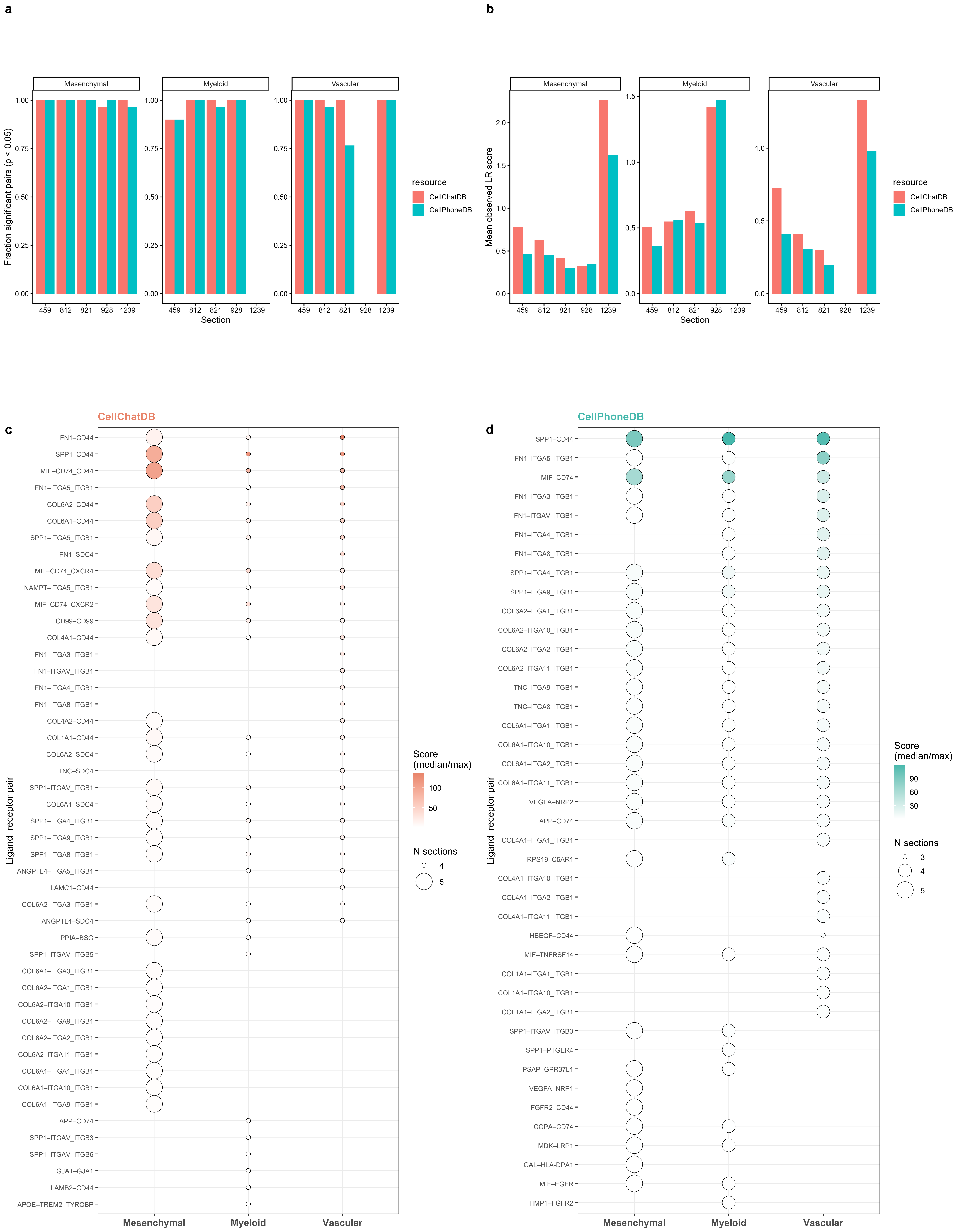
a
Mesenchymal
Myeloid
Vascular
Fraction significant pairs (p < 0.05)
0.00
0.25
0.50
0.75
1.00
459
812
821
928
1239
Section
resource
CellChatDB
CellPhoneDB
b
Mesenchymal
Myeloid
Vascular
Mean observed LR score
0.0
0.5
1.0
1.5
2.0
459
812
821
928
1239
Section
resource
CellChatDB
CellPhoneDB
CellChatDB
c
FN1–CD44
SPP1–CD44
MIF–CD74_CD44
FN1–ITGA5_ITGB1
COL6A2–CD44
COL6A1–CD44
SPP1–ITGA5_ITGB1
FN1–SDC4
MIF–CD74_CXCR4
NAMPT–ITGA5_ITGB1
MIF–CD74_CXCR2
CD99–CD99
COL4A1–CD44
FN1–ITGA3_ITGB1
FN1–ITGAV_ITGB1
FN1–ITGA4_ITGB1
FN1–ITGA8_ITGB1
COL4A2–CD44
COL1A1–CD44
COL6A2–SDC4
TNC–SDC4
SPP1–ITGAV_ITGB1
COL6A1–SDC4
SPP1–ITGA4_ITGB1
SPP1–ITGA9_ITGB1
SPP1–ITGA8_ITGB1
ANGPTL4–ITGA5_ITGB1
LAMC1–CD44
COL6A2–ITGA3_ITGB1
ANGPTL4–SDC4
PPIA–BSG
SPP1–ITGAV_ITGB5
COL6A1–ITGA3_ITGB1
COL6A2–ITGA1_ITGB1
COL6A2–ITGA10_ITGB1
COL6A2–ITGA9_ITGB1
COL6A2–ITGA2_ITGB1
COL6A2–ITGA11_ITGB1
COL6A1–ITGA1_ITGB1
COL6A1–ITGA10_ITGB1
COL6A1–ITGA9_ITGB1
APP–CD74
SPP1–ITGAV_ITGB3
SPP1–ITGAV_ITGB6
GJA1–GJA1
LAMB2–CD44
APOE–TREM2_TYROBP
Ligand–receptor pair
Mesenchymal
Myeloid
Vascular
Score
(median/max)
100
50
N sections
4
5
CellPhoneDB
d
SPP1–CD44
FN1–ITGA5_ITGB1
MIF–CD74
FN1–ITGA3_ITGB1
FN1–ITGAV_ITGB1
FN1–ITGA4_ITGB1
FN1–ITGA8_ITGB1
SPP1–ITGA4_ITGB1
SPP1–ITGA9_ITGB1
COL6A2–ITGA1_ITGB1
COL6A2–ITGA10_ITGB1
COL6A2–ITGA2_ITGB1
COL6A2–ITGA11_ITGB1
TNC–ITGA9_ITGB1
TNC–ITGA8_ITGB1
COL6A1–ITGA1_ITGB1
COL6A1–ITGA10_ITGB1
COL6A1–ITGA2_ITGB1
COL6A1–ITGA11_ITGB1
VEGFA–NRP2
APP–CD74
COL4A1–ITGA1_ITGB1
RPS19–C5AR1
COL4A1–ITGA10_ITGB1
COL4A1–ITGA2_ITGB1
COL4A1–ITGA11_ITGB1
HBEGF–CD44
MIF–TNFRSF14
COL1A1–ITGA1_ITGB1
COL1A1–ITGA10_ITGB1
COL1A1–ITGA2_ITGB1
SPP1–ITGAV_ITGB3
SPP1–PTGER4
PSAP–GPR37L1
VEGFA–NRP1
FGFR2–CD44
COPA–CD74
MDK–LRP1
GAL–HLA-DPA1
MIF–EGFR
TIMP1–FGFR2
Ligand–receptor pair
Mesenchymal
Myeloid
Vascular
Score
(median/max)
90
60
30
N sections
3
4
5

**Fig. S2: Statistical robustness and full pair coverage of spatial ligand-receptor scoring across databases and sections.**

**(a)** Fraction of candidate ligand-receptor pairs reaching statistical significance ($p < 0.05$, permutation test, $n = 1000$ permutations) per section and tumor zone (Mesenchymal, Myeloid, Vascular), stratified by database (CellChatDB, salmon; CellPhoneDB, teal). Across most section-zone combinations, >90% of scored pairs exceeded the significance threshold in both databases. Missing bars indicate sections where insufficient zone spots precluded scoring for that zone.

**(b)** Mean observed LR score per section and zone, stratified by database, providing a measure of overall interaction signal strength independent of statistical significance. Section 1239 shows the highest mean observed score in the Mesenchymal zone (CellChatDB).

**(c, d)** Full top 30 ligand-receptor pair dot plots for CellChatDB (c) and CellPhoneDB (d) independently, prior to cross-database filtering. Dot size represents the number of sections in which the pair was detected; dot fill intensity represents the aggregated score (median for Mesenchymal and Myeloid zones; maximum for the Vascular zone).

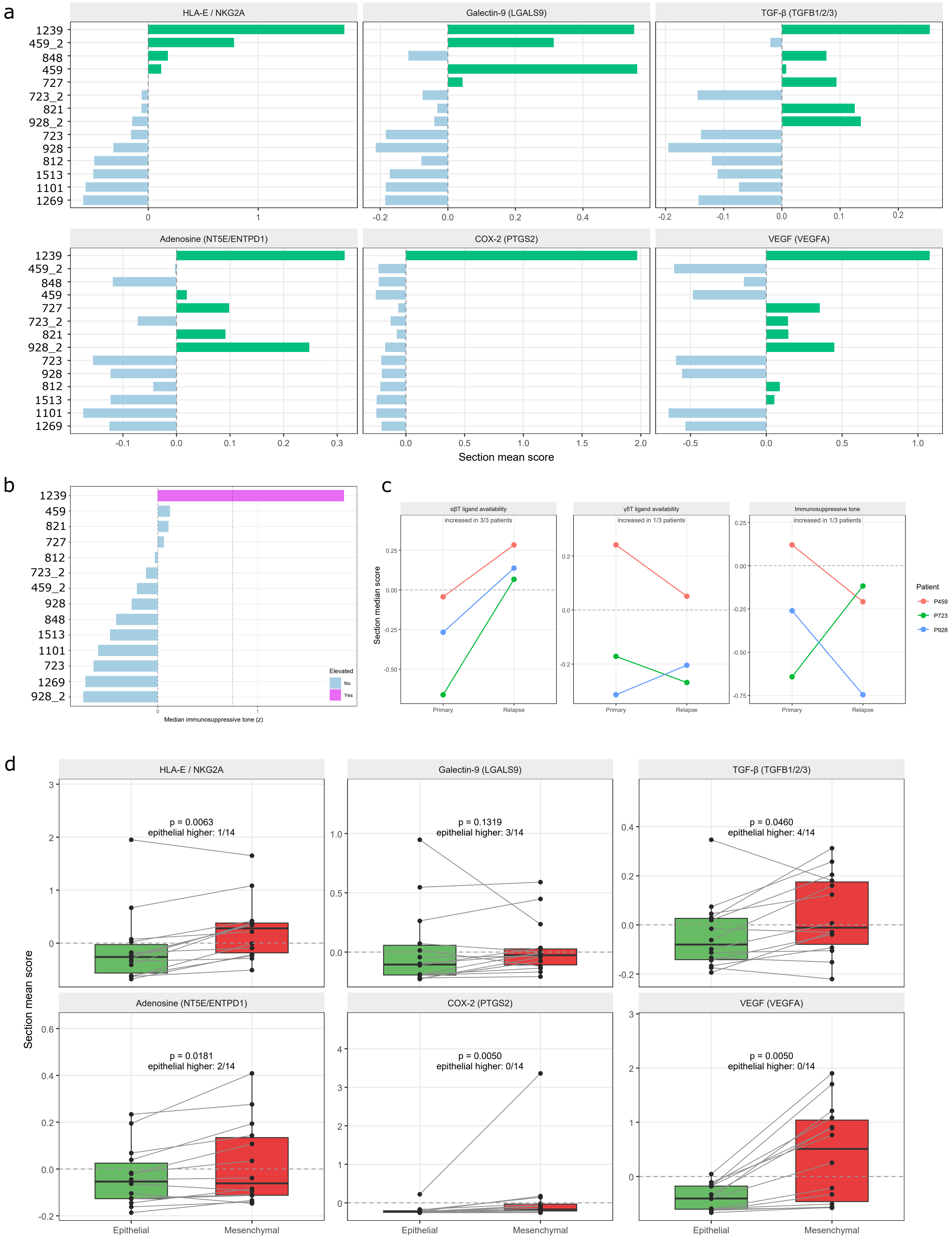
a
HLA-E / NKG2A
Galectin-9 (LGALS9)
TGF-β (TGFB1/2/3)
Adenosine (NT5E/ENTPD1)
COX-2 (PTGS2)
VEGF (VEGFA)
1239
459_2
848
459
727
723_2
821
928_2
723
928
812
1513
1101
1269
Section mean score
b
1239
459
821
727
812
723_2
459_2
928
848
1513
1101
723
1269
928_2
Elevated
No
Yes
Median immunosuppressive tone (z)
c
αβT ligand availability
increased in 3/3 patients
γδT ligand availability
increased in 1/3 patients
Immunosuppressive tone
increased in 1/3 patients
Section median score
Primary
Relapse
Patient
P459
P723
P928
d
HLA-E / NKG2A
p = 0.0063
epithelial higher: 1/14
Galectin-9 (LGALS9)
p = 0.1319
epithelial higher: 3/14
TGF-β (TGFB1/2/3)
p = 0.0460
epithelial higher: 4/14
Adenosine (NT5E/ENTPD1)
p = 0.0181
epithelial higher: 2/14
COX-2 (PTGS2)
p = 0.0050
epithelial higher: 0/14
VEGF (VEGFA)
p = 0.0050
epithelial higher: 0/14
Section mean score
Epithelial
Mesenchymal

**Fig. S3. Immunosuppressive mechanism axes and their relationship to tumor compartment, section, and relapse.**

**(a)** Immunosuppressive mechanism axes across the 14 ependymoma sections. Each bar shows the section's mean score for one of six mechanisms: HLA-E/NKG2A, galectin-9 (LGALS9), TGF-β (TGFB1/2/3), adenosine (NT5E/ENTPD1), COX-2 (PTGS2), and VEGF (VEGFA). Mechanisms are independent by design and are weighted equally in the immunosuppressive composite; axes are summarized as means rather than medians because for sparsely detected axes the section median reduces to zero on the z scale.

**(b)** Median immunosuppressive tone across the 14 sections. Section 1239 alone exceeds the elevated-tone threshold ($z = 0.75$, dotted line) used to define the Suppressed opportunity category; scores are z-scaled within the cohort and are therefore relative to these 14 sections.

**(c)** Matched primary versus relapse sections for the three opportunity components (αβT ligand availability, γδT ligand availability, immunosuppressive tone), shown separately rather than as fused net scores because a change in a net score cannot distinguish altered ligand availability from altered immunosuppressive tone. Each line connects the primary and relapse section from one patient ($n = 3$). No significance test is reported: with three matched pairs the smallest attainable p-value is 0.25.

**(d)** Immunosuppressive mechanism axes stratified by tumor compartment (epithelial versus mesenchymal), pooled across all 14 Visium sections. Boxplots show section-level mean scores for each mechanism; individual points represent sections, and lines connect matched epithelial and mesenchymal values from the same section. Paired Wilcoxon signed-rank tests compare compartments, with Benjamini–Hochberg-adjusted p-values reported within the six-mechanism family, alongside the number of sections in which the epithelial compartment scored higher. VEGF (VEGFA) and COX-2 (PTGS2) also contribute to the mesenchymal zone program used to define compartments, so those two comparisons are not fully independent of the compartment definition.
